# p75 Neurotrophin Receptor Regulates the Timing of the Maturation of Cortical Parvalbumin Cell Connectivity and Promotes Ocular Dominance Plasticity in Adult Visual Cortex

**DOI:** 10.1101/392159

**Authors:** Elie Baho, Bidisha Chattopadhyaya, Marisol Lavertu-Jolin, Raffaele Mazziotti, Patricia N Awad, Pegah Chehrazi, Marianne Groleau, Celine Jahannault-Talignani, Nathalie T. Sanon, Elvire Vaucher, Fabrice Ango, Tommaso Pizzorusso, Laura Baroncelli, Graziella Di Cristo

## Abstract

By virtue of their extensive axonal arborisation and perisomatic synaptic targeting, cortical inhibitory Parvalbumin (PV) cells strongly regulate principal cell output and plasticity. An interesting aspect of PV cell connectivity is its prolonged maturation time course, which is completed only by end of adolescence. The p75 neurotrophin receptor (p75NTR) regulates a wide range of cellular function, including apoptosis, neuronal process remodeling and synaptic plasticity, however its role on cortical circuit development is still not well understood, mainly because localizing p75NTR expression with cellular and temporal resolution has, so far, been challenging.

Using RNAscope and a modified version of the Proximity Ligation Assay (PLA), we show that p75NTR mRNA and protein are expressed in cortical PV cells in the postnatal and adult brain. Further, p75NTR expression in PV cells decreases between postnatal day (P)14 and P26, at a time when PV cell synapse numbers increase dramatically. Conditional knockout of p75NTR in single PV neurons in cortical organotypic cultures and in PV cell networks *in vivo* leads to precocious formation of PV cell perisomatic innervation and perineural nets around PV cell somata, suggesting that p75NTR expression controls the timing of the maturation of PV cell connectivity in the adolescent cortex.

Remarkably, we found that p75NTR is still expressed, albeit at low level, in PV cells in adult cortex. Interestingly, activation of p75NTR onto PV cells in adult visual cortex *in vivo* is sufficient to destabilize their connectivity and to reintroduce juvenile-like cortical plasticity following monocular deprivation. Altogether, our results show that p75NTR activation dynamically regulates PV cell connectivity, and represents a novel tool to foster brain plasticity in adults.

## INTRODUCTION

Within the forebrain, GABAergic (γ-aminobutyric acid producing) interneurons possess the largest diversity in morphology, connectivity, and physiological properties. A fascinating hypothesis is that different interneurons play partially distinct roles in neural circuit function and behavior. The large majority of cortical parvalbumin (PV)-positive interneurons specifically target the soma and proximal dendrites of pyramidal cells, and have been implicated in synchronizing the firing of neuronal populations and generating gamma oscillations^1–3^, which are important for perception, selective attention, working memory and cognitive control in humans and rodents^4–7^. Importantly, PV cells are also involved in experience-dependent refinement of cortical circuits during postnatal development, or critical period plasticity. Indeed, many studies on the visual cortex have demonstrated that the timing of critical period plasticity is set by PV cell maturation^8–12^. Furthermore, reducing GABAergic inhibition has been shown to partly restore juvenile-like plasticity in adult visual cortex^13,14^. However, whether alteration of PV cell connectivity and function is a necessary step for this effect is still unclear.

PV cell function relies on its pattern of connectivity: they innervate hundreds of postsynaptic targets with multiple synapses clustered around the cell body and proximal dendrites. In addition, PV cell connectivity has a prolonged developmental time course, plateauing towards the end of adolescence^15^. Recent studies have started to explore the molecular players underlying this unique innervation pattern, either in a cell-autonomous^12,16–18^ or cell non-autonomous fashion^11,19,20^. Conversely, the involvement of inhibitory mechanisms in the establishment of PV cell connectivity during development is less clear. In addition, it is unknown whether similar inhibitory molecular mechanisms could be recruited in the adult brain to reduce PV cell connectivity, and in parallel, increase experience-dependent plasticity.

The neurotrophin receptor p75NTR is a multifunctional receptor modulating several critical steps in nervous system development and function, from apoptosis and neuron morphology development to synaptic plasticity^21^. In particular, it has been shown that p75NTR interaction with the precursor form of BDNF, proBDNF, or with its prodomain alone, induces growth cone collapse and dendritic spine remodeling in hippocampal excitatory neurons^22,23^ and alterations in this process may lead to long term cognitive dysfunctions^23^. Due to the difficulty of pinpointing p75NTR localisation in cortical tissue with temporal and single cell resolution, whether and how p75NTR plays a role on cortical GABAergic circuit development is not well understood.

Using RNAscope and Proximity Ligation Essay, here, we show that cortical PV cells expressed p75NTR and that its expression level decreased during the maturation phase of PV cell connectivity. Conditional knockout of p75NTR in single PV cells promoted the formation of their perisomatic innervation in cortical organotypic cultures. This effect was mimicked by transfection of a p75NTR dominant negative form in wild-type PV cells and was rescued by expression of p75NTR in p75NTR^-/-^ PV cells. Conversely, increasing p75NTR signaling strongly reduced PV cell connectivity, both in young and mature organotypic cortical cultures. Further, conditional knockout of p75NTR in GABAergic cells derived from the medial ganglionic eminence promoted the precocious formation of PV cell perisomatic innervation and perineural nets (PNN) around PV cell somata *in vivo*. These data suggest that p75NTR expression modulates the timing of the maturation of PV cell connectivity in the adolescent cortex. Finally, we observed that p75NTR activation in PV cells destabilized their innervation, dramatically reduced perineural net density and intensity and promoted ocular dominance plasticity in adult visual cortex. All together, these data suggest a novel, powerful role for p75NTR-mediated signaling in modulating PV cell connectivity, both during development and in adulthood.

## RESULTS

### Cortical PV cells express p75NTR during development and in the adult brain

The temporal and cellular localization of p75NTR in cortical neurons has been an object of debate and discrepancy^23,24^, likely due to low protein expression levels, especially in the adult brain, and suboptimal antibody sensitivity. To overcome these technical challenges, we used two novel experimental strategies. First, we used RNAscope multiplex fluorescent *in situ* hybridization (Advanced Cell Diagnostics), a novel RNA *in situ* hybridization technology, that allows single-molecule detection^25^, to detect both PV and p75NTR mRNA in brain slices (Fig. 1a, b; Supplementary Fig. 1). Importantly, we found cortical neurons co-expressing both PV and p75NTR (Fig. 1a, b), in contrast to basal ganglia wherein p75NTR and PV were expressed by clearly non-overlapping populations (Supplementary Fig. 1e). Secondly, we used a modified version of the proximity ligation essay (PLA) as described in Telley et al.^26^, coupled with PV immunolabeling. PLA is a very sensitive technique of amplification utilized to detect low level of protein expression or protein-protein interaction in tissues, using which we observed unprecedented clear and definite signal for p75NTR in PV cell somata and putative boutons in visual cortex of adult mice (P60) (Fig. 1c). To control for PLA signal specificity, we crossed p75NTR^lox/lox^ mice with mice expressing Cre recombinase under the control of the PV promoter (PV_Cre)^36^ and compared PLA-p75NTR labeling in p75NTR^lox/lox^ vs. PV_Cre;p75NTR^lox/lox^ littermates (Fig. 1c-f). We found that p75NTR signal was dramatically reduced in PV cells in the conditional knockout mice (Fig. 1e; unpaired t-test, p = 0.0006), demonstrating the specificity of our approach. Surprisingly, we also observed that the total p75NTR signal showed a ∼60% reduction in PV_Cre;p75NTR^lox/lox^ mice compared to control littermates (Fig. 1f; unpaired t-test, p = 0.004), suggesting that a large proportion of p75NTR protein was expressed by PV cells in the adult visual cortex.

**Figure 1.**
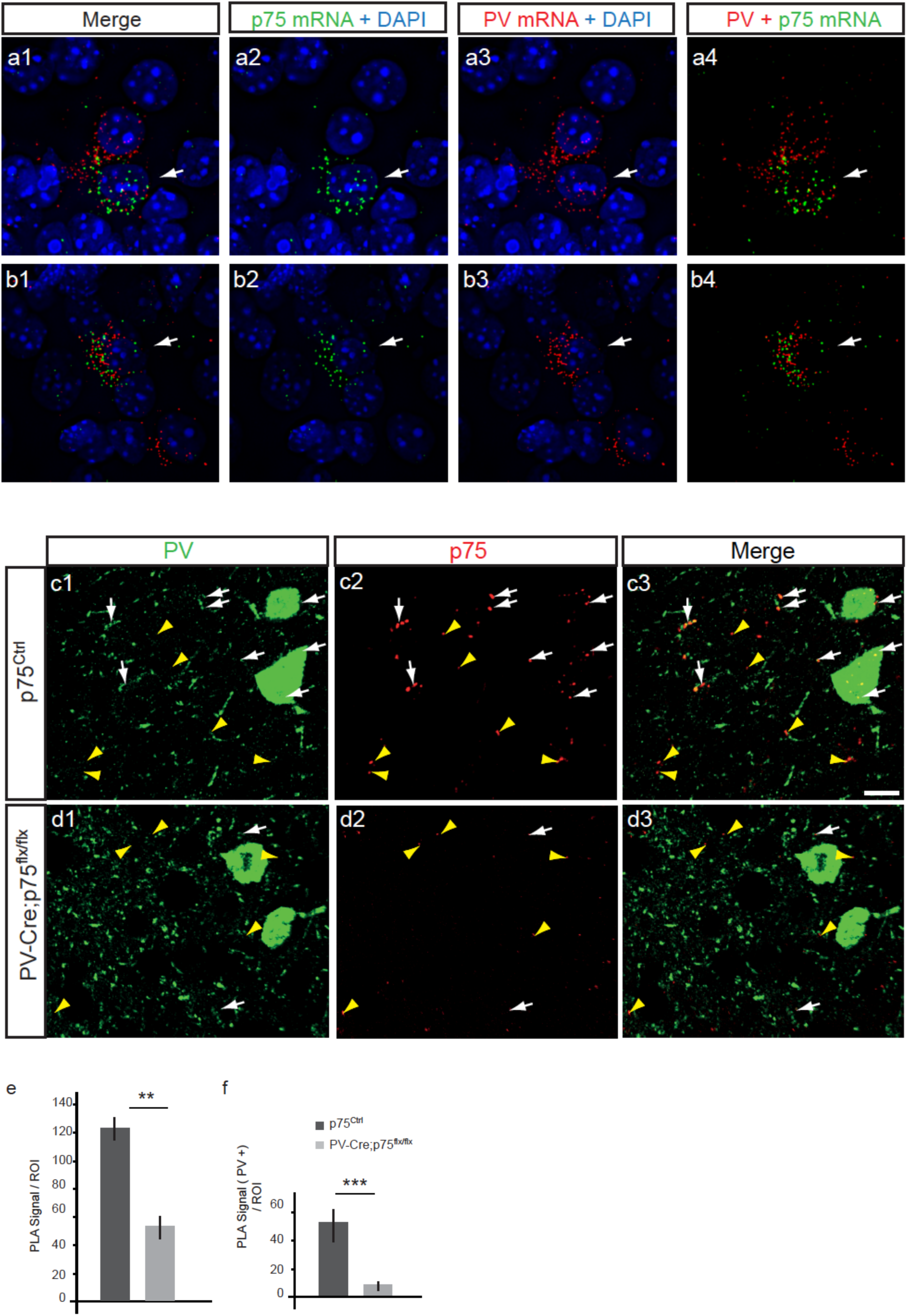
A subset of PV cells express p75NTR mRNA and proteins in adult cortex. (a, b). Images from coronal brain section hybridized with PV and p75 probes using fluorescent multiplex RNAscope technology. p75 mRNA (a2, b2; green dots) can be detected in cells expressing PV mRNA (a3, b3; red dots). White arrows point to p75 and PV signals around the same nucleus, identified by DAPI staining (blue). (c, d) Cortical slices from p75NTR^flx/flx^ (c) or PV-Cre;p75NTR^flx/flx^ (c) co-immunostained with PV (c1, d1; green) and p75NTR using PLA (c2, d2; Red dots). White arrows point to PLA signals that colocalize with PV signals (c1-3, d1-3). Note that p75NTR signal can be observed in PV cell boutons. Yellow arrowheads show PLA signals without PV colocolization (c1-3, d1-3). Scale bar: 10µm (e) Quantification of PLA signal reveal a significant reduction of total PLA signals per ROI in PV-Cre;p75NTR^flx/flx^ as compared to wild-type littermates. t-test, p=0.004. (f) Further, PLA signals that co-localized with PV labeling decrease significantly in PV-Cre;p75NTR^flx/flx^ as compared to wild-type littermates. t-test, p=0.0006.

Next, we asked whether p75NTR protein expression in PV cells was developmentally regulated, in visual cortex. We found that p75NTR expression in PV cells was significantly reduced between P14 and P26 (Fig. 2a-c, unpaired t-test with Welch’s correction, p < 0.001). In comparison to its expression in adult visual cortex, we observed similar localization pattern of p75NTR protein in PV cell somata and in putative perisomatic synapses at both P14 and P26. Overall, these data suggest that cortical PV cells express p75NTR and that this expression is developmentally regulated.

**Figure 2.**
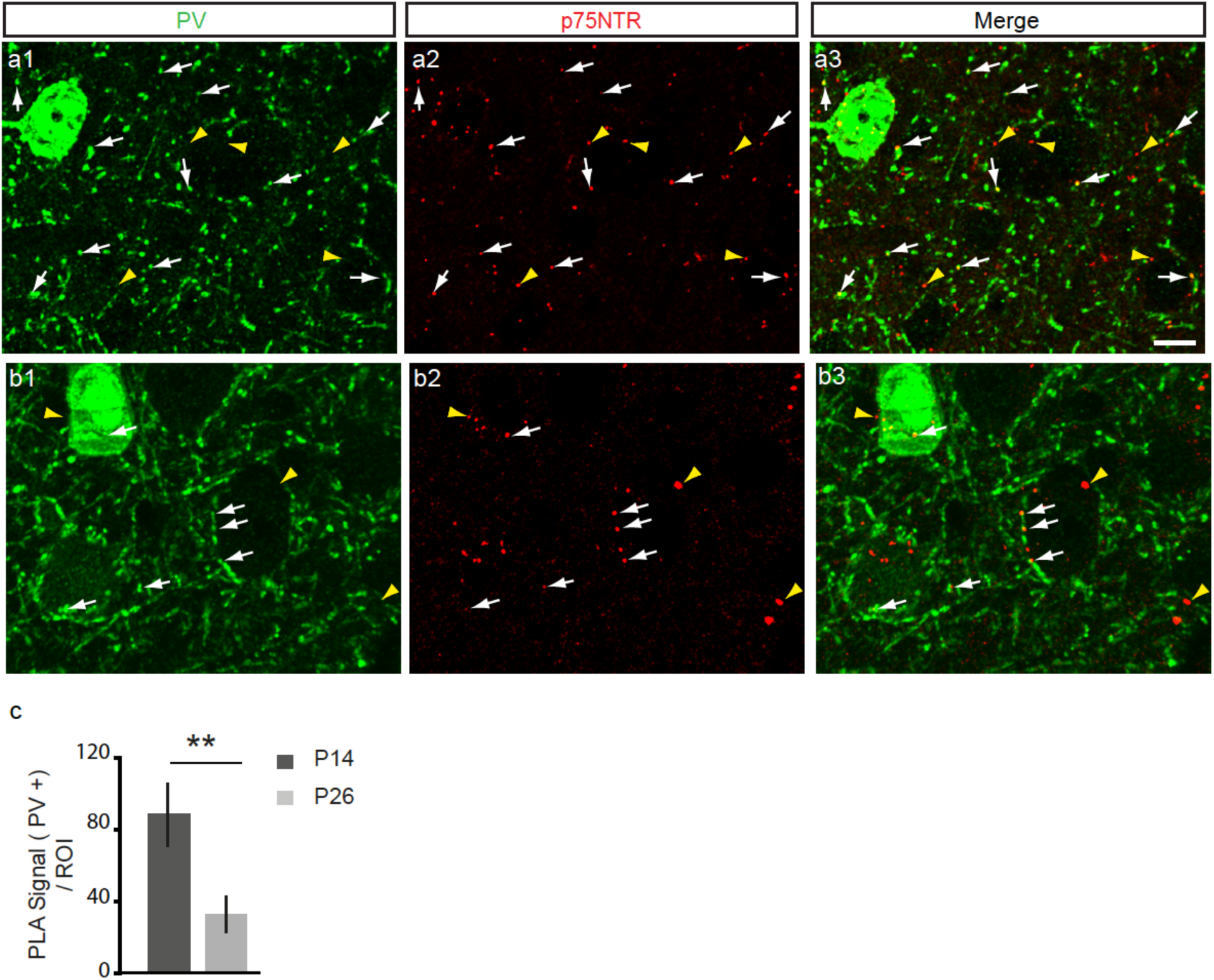
p75NTR expression in cortical PV cells decreases during the first postnatal month. (a, b) Cortical slices from P14 (a) and P26 (b) wild-type mice immunostained with parvalbumin (PV) to label PV cells (a1, b1; green) and PLA-mediated labeling for p75NTR (a2, b2; red dots, henceforth indicated in figures as p75). White arrows point to PLA signals that co-localize with PV signals (a1-3, b1-3). Yellow arrowheads show PLA signals without PV co-locolization (a1-3, b1-3). Note that at both ages, p75NTR signal can be found in putative PV cell boutons. Scale bar: 10µm. (c) Quantification of p75NTR PLA intensity in PV cells at different postnatal ages shows a significantly decline of p75NTR signal in PV cells and boutons between P14 and P26 (unpaired t-test with Welch’s correction, p<0.001). n=2 animals for each age point.

### p75NTR downregulation during the first postnatal weeks induce the formation of exuberant PV cell innervation in cortical organotypic cultures

Since the developmental down-regulation of p75NTR was inversely correlated with the maturation of PV cell innervation during the same time period in visual cortex^15^, we hypothesized that higher p75NTR levels may hinder the formation of PV cell synapses. To test this hypothesis, we used cortical organotypic cultures where we could label and manipulate isolated PV cells by driving GFP and/or Cre expression with a previously characterized promoter (P_G67_; ^10,15–17^). In organotypic cultures, PV cells start out with very sparse and simple axons, which develop into complex, highly branched arbors in the subsequent 3 weeks with a time course similar to that observed *in vivo*^15^.

In the postnatal cortex, p75NTR is not expressed exclusively by PV cells (Fig. 2; and ^27^), thus to investigate whether p75NTR in PV cells plays a role in their maturation, we knocked-down p75NTR expression selectively in PV cells by transfecting P_G67__Cre/GFP in organotypic cultures from p75NTR^lox/lox^ mice^28^ to generate p75NTR^-/-^ PV cells in an otherwise wild-type background (Fig. 3). PV cells were transfected with P_G67__Cre/GFP from Equivalent Postnatal day (EP)10 (cultures prepared at P4 + 6 days *in vitro*) and fixed at EP18. p75NTR^-/-^ PV cells contacted more pyramidal cells and formed more axonal branching and perisomatic boutons as compared to age-matched control p75NTR^lox/lox^ PV cells, which were transfected with P_G67__GFP alone (Fig. 3a, b, c, e; perisomatic bouton density, unpaired t-test, p < 0.001; percentage of innervation, unpaired t-test, p < 0.001). p75NTR reduction in single PV cells during the peak of perisomatic bouton proliferation (EP16-24) also increased bouton density and terminal branching without increasing the percentage of contacted cells (Fig. 3c-e; perisomatic bouton density, Mann Whitney Rank Sum test, p = 0.002; percentage of innervation, unpaired t-test, p = 0.166). These data suggest that p75NTR expression constrains the maturation of PV cell innervation in a cell-autonomous manner.

**Figure 3.**
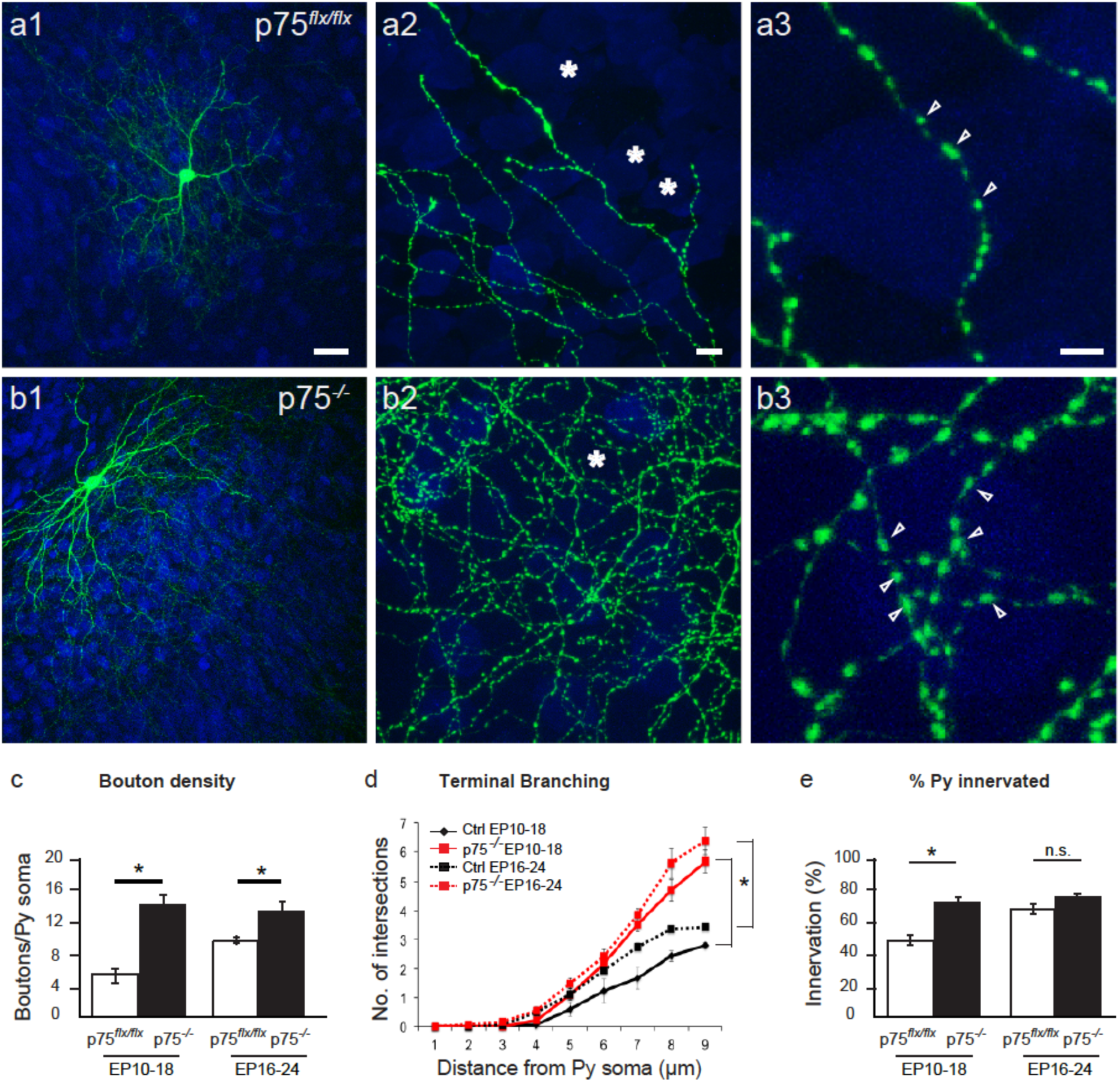
Cre-mediated inactivation of p75NTR in single PV cells induces the formation of more complex innervations. (a) Control PV cell transfected with P_g67_-GFP (Ctrl, green) in EP18 organotypic cultures from *p75^flx/flx^* mice. (b) PV cells transfected with P_G67_-Cre/GFP from EP10-18 (p75^-/-^ PV cells) shows perisomatic innervation characterized by multiple terminal axonal branches (b2) bearing numerous clustered boutons (b3; arrowheads) around pyramidal cell somata (NeuN immunostaining, blue). Stars indicate pyramidal cells somata that are not innervated. (a3) and (b3) are from regions in (a2) and (b2). Scale bar, a1, b1: 50µm; a2, b2: 5µm; a3, b3: 3µm. Perisomatic boutons density (c), terminal branching (d) and percentage of innervated cells (e) of p75^flx/flx^ and p75^-/-^ PV cells transfected at EP10-18 or EP16-24 (c) EP10-18: unpaired t-test, p<0.001, EP16-24: Mann Whitney test, p=0.002. (d) EP10-18: Mann Whitney test p<0.001, EP16-24: unpaired t-test, p<0.001. (e) EP10-18: unpaired t-test<0.001, EP16-24: unpaired t-test, p=0.166. EP10-18; n = 8 p75^-/-^ PV cells, n = 7 p75^flx/flx^ PV cells. EP16-24; n = 6 p75^-/-^ PV cells, n = 6 p75^flx/flx^ PV cells.

To further test this hypothesis, we investigated whether transfecting wild-type PV cells with a mutant form of p75NTR lacking the intracellular death domain (p75ΔDD^21,29^) could affect their innervation (Fig. 4a, b). Since the death domain is critical for protein-protein interactions, we reasoned that p75ΔDD would act as a dominant negative. PV cells transfected with p75ΔDD showed more complex perisomatic innervation (Fig. 4a, b, e; perisomatic bouton density, one-way ANOVA, p < 0.0001, *post hoc* Tukey’s test p75 ΔDD vs p75^lox/lox^, p = 0.0002), which was indistinguishable from those formed by p75NTR^-/-^ PV cells (Fig. 4 b, c, e; perisomatic bouton density, p75NTR^-/-^ vs p75 ΔDD, *post hoc* Tukey’s test, p = 0.1314).

**Figure 4.**
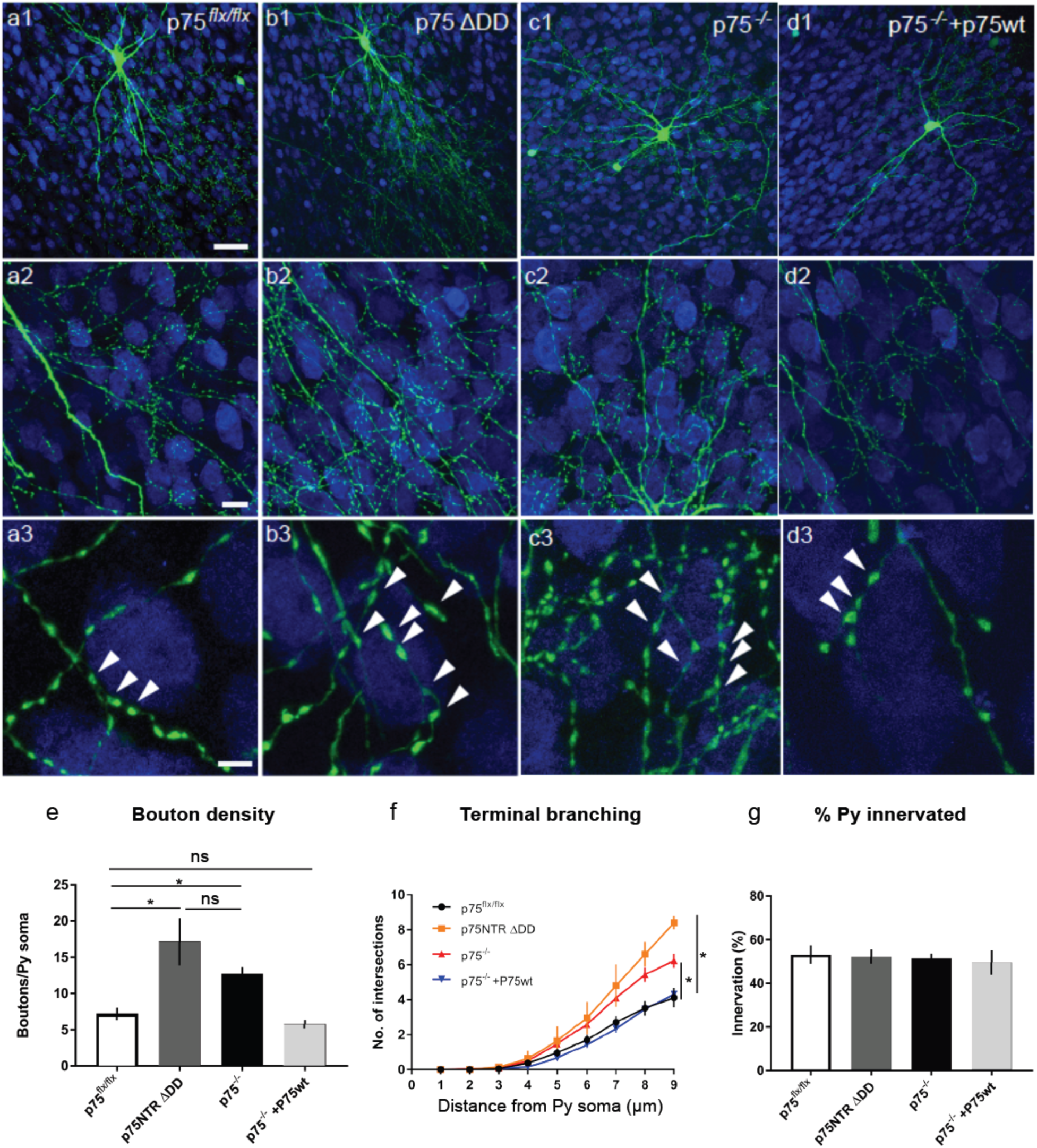
p75NTRΔDD mimics, while p75NTRwt rescues the innervation phenotype of p75NTR^-/-^ PV cells. (a) Control PV cell transfected with P_g67_-GFP (Ctrl, green) in EP24 organotypic cultures from p75^flx/flx^ mice. (b) PV cells transfected with P_G67_-GFP/ p75NTRΔDD from EP16-24 (p75^-/-^ p75ΔDD PV cells) shows more complex perisomatic innervation characterized by multiple terminal axonal branches (c2) bearing numerous clustered boutons (c3; arrowheads) around pyramidal cell somata (NeuN immunostaining, blue). (c) PV cells transfected with P_G67_-Cre/GFP (p75^-/-^ PV cells) resemble p75ΔDD PV cells. (d) p75^-/-^ PV cells transfected with p75NTR cDNA (p75^-/-^ + p75wt PV cells) are indistinguishable from control PV cells. a3, b3, c3, d3 are from regions in a2, b2, c2, d3. Scale bar, a1, b1: 50µm; a2, b2: 10µm; a3, b3: 5µm. Perisomatic boutons density (e), terminal branching (f) and percentage of innervated cells (g) (e) One way Anova with *post hoc* Tukey’s test. p75^flx/flx^ vs p75ΔDD PV cells, p=0.0002; p75^flx/flx^ vs p75^-/-^ PV cells, p=0.0141; p75^flx/flx^ vs p75^-/-^ + p75wt PV cells, p=0.8533; p75ΔDD vs p75^-/-^ PV cells, p=0.1314. (f) One way Anova with *post hoc* Tukey’s test, p<0.001 at 7, 8 and 9 μm from pyramidal (Py) soma center. (g) One way Anova, p>0.05. PV cells: n = 9 p75^flx/flx^, n= 5 p75ΔDD, n = 9 p75^-/-^ PV cells, n = 7 p75^-/-^ + p75wt.

It is conceivable that Cre-mediated removal of exons 4-6 in *p75NTR* might also delete intronic sequences that are important for PV cell synaptic development. To confirm that the deletion of p75NTR was indeed responsible for the exuberant perisomatic innervation of p75NTR^-/-^ PV cells, we performed a rescue experiment by introducing p75NTR cDNA in p75NTR^-/-^ PV cells. In particular, we co-transfected PV cells in organotypic cultures prepared from p75NTR^lox/lox^ mice with either P_G67__Cre/GFP (to generate p75NTR^-/-^ PV cells) or P_G67__Cre/GFP/p75wt (to re-express p75NTR in p75NTR^-/-^ PV cells). The perisomatic innervation formed by reintroduction of p75NTR in p75NTR^-/-^ PV cells was not significantly different from those formed by wild-type cells (Fig. 4a, d, e; *post hoc* Tukey’s test, p = 0.8533).

All together, these data demonstrate that p75NTR expression in cortical PV cells regulates the maturation of their connectivity, by constraining the formation of their perisomatic innervation.

### Increased p75NTR activation inhibits the formation of PV cell innervation in cortical organotypic cultures

p75NTR-mediated signaling can be strongly activated by proneurotrophins and their prodomain^22,30–33^. Activation of p75NTR by proBDNF has been shown to reduce excitatory synapse density in hippocampal pyramidal neurons^34,35^, to promote excitatory synapse elimination in the postnatal visual cortex^36^ and at the developing neuromuscular junction^37^. To investigate whether proBDNF affects the development of inhibitory PV cell connectivity, we treated developing organotypic cultures either with a wild-type recombinant form (wt-proBDNF, 10ng/ml) or a cleavage-resistant mutant form of proBDNF (mut-proBDNF; 10ng/ml) from EP18-24. wt-proBDNF did not significantly affect PV cell perisomatic bouton number (Supplementary Fig. 2a, b, e), but induced a significant increase of the terminal axonal branching complexity formed by PV cells around their targets (Supplementary Fig. 2f, One-way ANOVA with *post hoc* Tukey’s test, p < 0.05). It is likely that proBDNF was at least partially cleaved by extracellular plasmin and metalloproteases, thus affecting the local, relative level of mBDNF and proBDNF^38^. On the other hand, PV cells treated with mut-proBDNF contacted less than half of the pyramidal cells compared to age-matched controls, onto which they formed fewer boutons and terminal axonal branching (Supplementary Fig. 2 c, e-g; perisomatic bouton density, One-way ANOVA with *post hoc* Tukey’s test, Ctrl vs mut-proBDNF p = 0.0139; percentage of innervation, One-way ANOVA with *post hoc* Tukey’s test, Ctrl vs mut-proBDNF, p < 0.0001). This effect was not secondary to neuronal death because neuron density (based on NeuN immunostaining) was not altered compared to control or wt-proBDNF treated slices even after 6 days of treatment (Ctrl: 104 ± 13, wt-proBDNF: 174 ± 10 and mut-proBDNF: 135 ± 15 × 10^3^ pyramidal cells/mm^3^; n = 6 ctrl slices, n = 6 wt-proBDNF treated slices, n = 8 mut-proBDNF treated slices; One-way ANOVA, p > 0.05). To investigate whether the effects of mut-proBDNF on PV cell innervation were specifically mediated by p75NTR activation, we knocked-down p75NTR from single PV cells in organotypic cultures prepared from p75NTR^lox/lox^ mice and simultaneously treated them with mut-proBDNF. We found that p75NTR^-/-^ PV cells were insensitive to mut-proBDNF treatment; in fact, they formed significantly more complex innervations compared to both control and mut-proBDNF treated p75NTR^lox/lox^ PV cells (Supplementary Fig. 2 a, d, e-g; perisomatic bouton density, One-way ANOVA with *post hoc* Tukey’s test, Ctrl vs p75^-/-^ + mut-proBDNF, p < 0.0001). The results suggest that specific activation of p75NTR strongly inhibits the formation of PV cell innervation during postnatal development.

Several studies suggested that mature BDNF (mBDNF) and BDNF prodomain (pBDNF) are the most abundant moieties in the adult brain, while proBDNF is abundant during early development, in particular during the first postnatal month^22,31,32,34^. We thus asked whether altering endogenous levels of proBDNF and mBDNF affected the establishment of PV cell innervation in the first postnatal weeks. One of the molecular mechanisms responsible for the activity-dependent cleavage of proBDNF into mBDNF in the extracellular space is tissue plasminogen activator (tPA)-mediated activation of plasmin^38,39^. To alter tPA activity levels, we treated organotypic cultures with either PPACK (50 μM), a tPA-inactivating peptide, or tPA (0.6 µg/µl) from EP10-18, when PV cell axonal arborization and synaptic innervation are still quite immature^15^. Firstly, we sought to quantify whether and how endogenous mBDNF and proBDNF levels were affected by these treatments by western blot. While we confirmed the specificity of the anti-mBDNF antibody using brain lysates of BDNF KO mice (Supplementary Fig. 3a), we tested several commercial proBDNF antibodies, but, in our hands, they could still detect a 32kDa band in brain lysates from BDNF KO mice (see Methods for details on tested antibodies), thus we could only quantify mBDNF levels. As predicted, we found that treatment with PPACK reliably induced a significant reduction (Supplementary Fig. 3b), while tPA significantly increased, mBDNF protein level (Supplementary Fig. 3c), suggesting that tPA may indeed regulate extracellular level of mBDNF in this developmental time window. Consistent with this hypothesis, PV cells in PPACK-treated cultures showed simpler innervation fields (Supplementary Fig. 4a, b, e-g), while tPA addition drastically increased the complexity of PV cell axonal arborization compared to control, age-matched PV cells by increasing bouton density, terminal axonal branching and percentage of innervated targets (Supplementary Fig. 4 a, c, e-g).

Since mBDNF-mediated TrkB signaling is a potent regulator of GABAergic cell maturation^19,20,40^, it is possible that these effects might be solely due to alteration of mBDNF level, independently of p75NTR activation. First, we reasoned that, if this was indeed the case, then PPACK treatment should reduce perisomatic innervation formed by p75NTR^-/-^ PV cells compared to those formed by untreated p75NTR^-/-^ PV cells, since mBDNF level was reduced in presence of PPACK (Supplementary Fig. 4). However, similar to what we observed following the treatment with recombinant mut-proBDNF (Supplementary Fig. 2), the effects of PPACK on PV cell innervation were dependent upon the expression of p75NTR by PV cells. In fact, PPACK-treated p75NTR^-/-^ PV cells were indistinguishable from age-matched, untreated p75NTR^-/-^ PV cells (Supplementary Fig. 5; One-way ANOVA with *post hoc* Holm Sidak test, PPACK-treated p75NTR^-/-^ PV cells vs p75NTR^-/-^ PV cells, p > 0.1). Secondly, we reasoned that if the effects of tPA application on PV cell innervation was mediated by a decrease in proBDNF-mediated p75NTR signaling, then treatment with mut-proBDNF would reverse them. Supporting this prediction, we found that simultaneously treating organotypic cultures with tPA and mut-proBDNF rescued completely the effects of tPA-only application (Supplementary Fig. 4d, e-g).

In summary, these results suggest that p75NTR activation, possibly mediated by endogenous proBDNF, can strongly inhibit the formation of cortical PV cell innervation during the first postnatal weeks.

### p75NTR regulates the timing of the maturation of PV cell connectivity *in vivo*

Our results show that p75NTR expression in PV cells declines during the maturation phase of PV cell connectivity and that removing p75NTR is sufficient to promote, while activating p75NTR inhibits, the formation of PV cell innervation. We next asked whether p75NTR plays a role in the maturation of PV cell connectivity *in vivo*.

In PV_Cre mice, Cre expression is very specific to cortical PV cells, however it starts after P10 and does not plateau until weeks later^41^. Thus, to reduce p75NTR expression in PV cells before the peak of the maturation of PV cell connectivity, we generated Nkx2.1_Cre;p75NTR^lox/lox^ mice. Nkx2.1 is expressed in GABAergic precursors originating from the medial ganglionic eminence, which include PV and somatostatin expressing interneurons^42^. We quantified the putative perisomatic synapses formed by PV cells, identified by the juxtaposition of PV and gephyrin, a scaffolding protein present in the postsynaptic sites of GABAergic synapses, in the visual cortex of P14 Nkx2.1_Cre; p75NTR^lox/lox^ mice compared to their control littermates (Fig. 5a, b). Both the density of PV+gephyrin+ puncta and the percentage of perisomatic PV puncta showing gephyrin co-labelling were significantly increased (Fig. 5c, d; unpaired t-test, p = 0.04 for both graphs).

**Figure 5.**
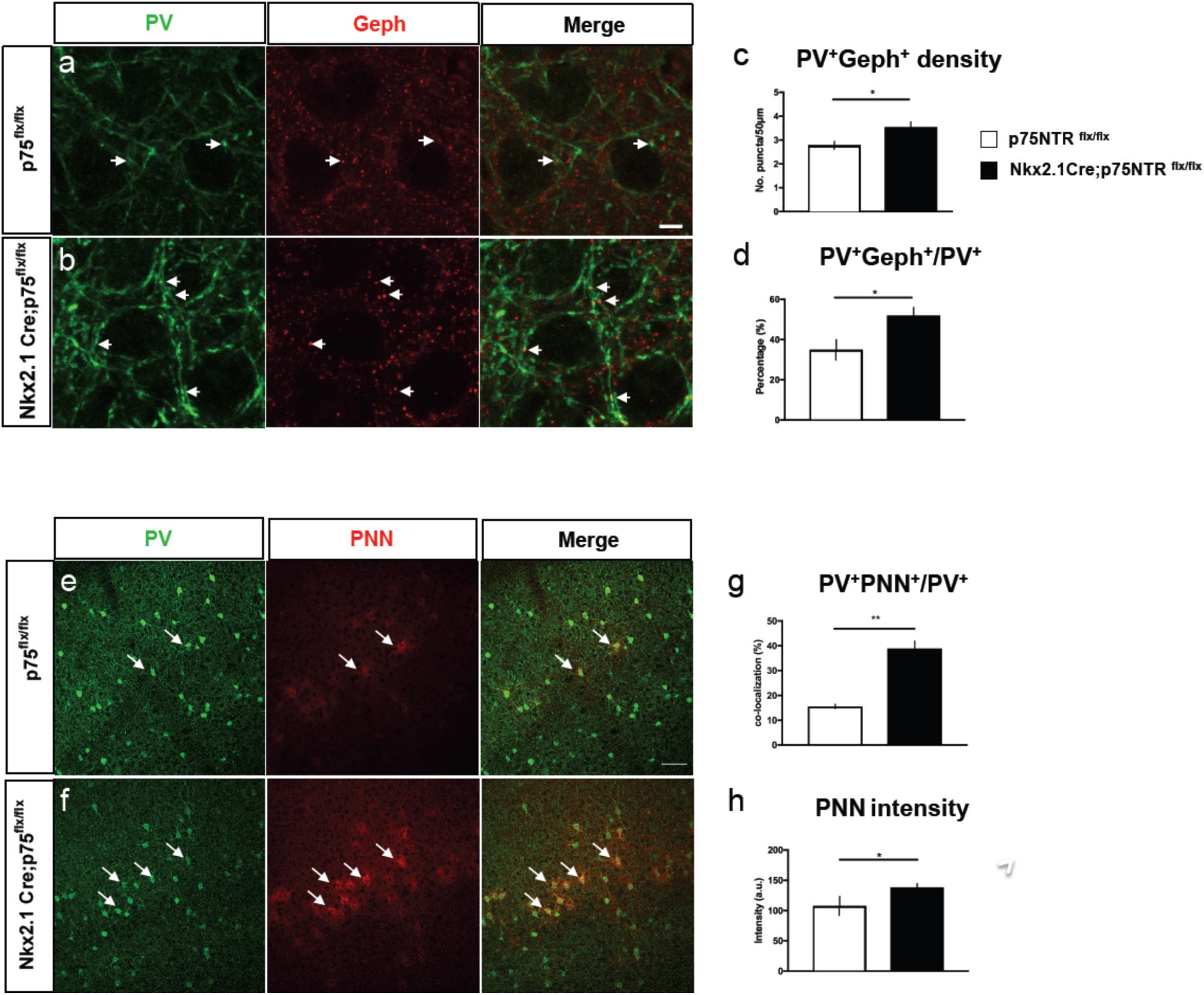
Cortical PV cells form more perisomatic boutons and are precociously enwrapped by PNN in Nkx2.1Cre; P75NTR^flx/flx^ mice. (a, b) Cortical slices from P14 p75NTR^flx/flx^ (a) or Nkx2.1-Cre;p75NTR^flx/flx^ (b) co-immunostained with PV (green) and gephryn (Geph, red). Arrows indicate examples of perisomatic PV+/Geph+ puncta. (c, d) Perisomatic PV+/Geph+ density (c) and percentage of PV+ puncta co-labeled with gephryn (d) are significantly increased in Nkx2.1Cre; p75NTR^flx/flx^ mice compared to control littermates. (c) Unpaired t-test, p=0.0407. (d) Unpaired t-test, p=0.0429. N=4 p75NTR^flx/flx^ and 6 Nkx2.1Cre; p75NTR^flx/flx^ mice. (e, f) Cortical slices from P18 p75NTR^flx/flx^ (e) or Nkx2.1-Cre;p75NTR^flx/flx^ (f) labeled with anti-PV antibody (green) and WFA, which stains perineural nets (PNN, red). Arrows indicate examples of PV+ somata enwrapped in PNN. (g, h) The proportion of PV somata surrounded by PNN (g) and mean PNN intensity (d) are significantly increased in Nkx2.1Cre; p75NTR^flx/flx^ mice compared to control littermates. (c) Unpaired t-test, p=0.0018. (d) Unpaired t-test, p=0.0343. N=3 mice for both genotpyes.

One important indication of PV cell maturation is the appearance of perineuronal nets (PNN), which enwrap the soma and primary dendrites of mature PV cells^43,44^. In Nkx2.1_Cre; p75NTR^lox/lox^, we observed a significant increase in both the number of PV cells that were encircled by PNN, as revealed by WFA staining (Fig. 5 e, f, g; unpaired t-test, p = 0.0018) and PNN immunofluorescence intensity around single PV cell somata (Fig. 5 h; unpaired t-test, p = 0.0343). Overall, these data demonstrate that p75NTR expression level regulates the timing of PV cell maturation *in vivo*.

### p75NTR activation destabilizes PV cell connectivity in adult brain

Our expression studies show that p75NTR is still expressed, albeit at a low level, in cortical PV cells in adult mice (Fig. 1c, e, f). We thus wondered whether activation of p75NTR might destabilize PV cell connectivity after it had reached maturity (around the 4^th^ postnatal week, both in cortical organotypic cultures and *in vivo*^15^). In cortical organotypic cultures, PV cells treated with mut-proBDNF from EP26-32, after PV cell innervation have plateaued, show a dramatic loss in both synaptic contacts and complexity of perisomatic innervation as compared to age-matched, control PV cells (Fig. 6a, c, d-f; One-way ANOVA, *post hoc* Holm-Sidak test, p < 0.05), while treatment with wt-proBDNF did not affect any of the analyzed parameters (Fig. 6b, d-f). Next, we asked whether treatment with mut-proBDNF could destabilize PV cell innervation in the adult brain *in vivo*. To address this question, we implanted osmotic minipumps releasing either mut-proBDNF (1 µg/ml, flow rate 0.5 µl/h) or vehicle solution in primary visual cortex in adult mice for 5 days (Fig. 7a). We found that in the cortices infused with mut-proBDNF (ipsilateral to the minipump), the density of perisomatic puncta immunopositive for the vesicular GABA transporter (vGAT, which labels presynaptic GABAergic terminals) or for PV was reduced as compared to those in the vehicle infused cortices (contralateral to the minipump) (Fig. 7b, d; Supplementary Fig. 9; ∼40% reduction for both PV+ and VGAT+ puncta/pyramidal soma in ipsicompared to contralateral cortex; unpaired t-test, p < 0.001).

**Figure 6.**
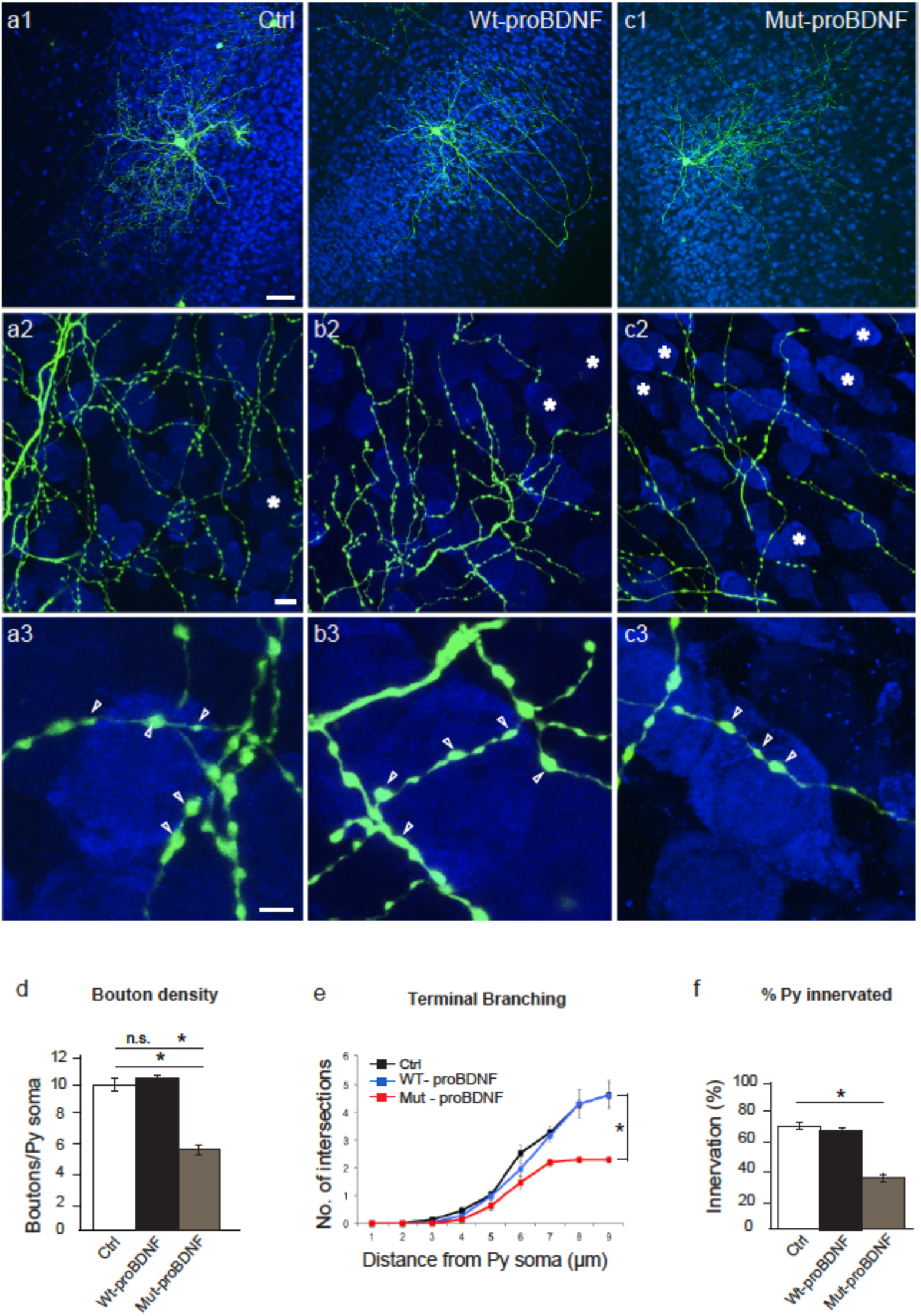
mut-proBDNF can destabilize PV cell innervation even after it has reached maturity. (a) Control PV cell (a1, Ctrl, green) at EP32 with exuberant innervation field characterized by extensive branching contacting the majority of potential targets, dense boutons along axons (a2), and terminal branches with prominent and clustered boutons (a3; arrowheads) around pyramidal cell somata (NeuN immunostaining, blue). (b) PV cell treated with wt-proBDNF from EP26-32 shows overall similar axon size (b1), percentage of potentially targeted neurons (B2) and perisomatic innervations (b3) as control, untreated PV cells. (c) PV cell treated with mut-proBDNF from EP26-32 shows a drastic reduction both in percentage of innervated cells (c2) and perisomatic innervation (c3). Stars indicate pyramidal cells somata that are not innervated. Scale bar, a1-c1: 50µm; a2-c2: 10µm; a3-c3: 5µm. (d) Perisomatic bouton density (e) terminal branching and (f) percentage of innervated cells of the three experimental groups. One-way Anova, *post hoc* Tukey test, p<0.0001 for Ctrl vs Mut-proBDNF and WT-proBDNF vs Mut-proBDNF for graphs in d-f. n = 9 Ctrl, n = 6 wt-proBDNF treated PV cells, n = 6 mut-proBDNF treated PV cells.

**Figure 7.**
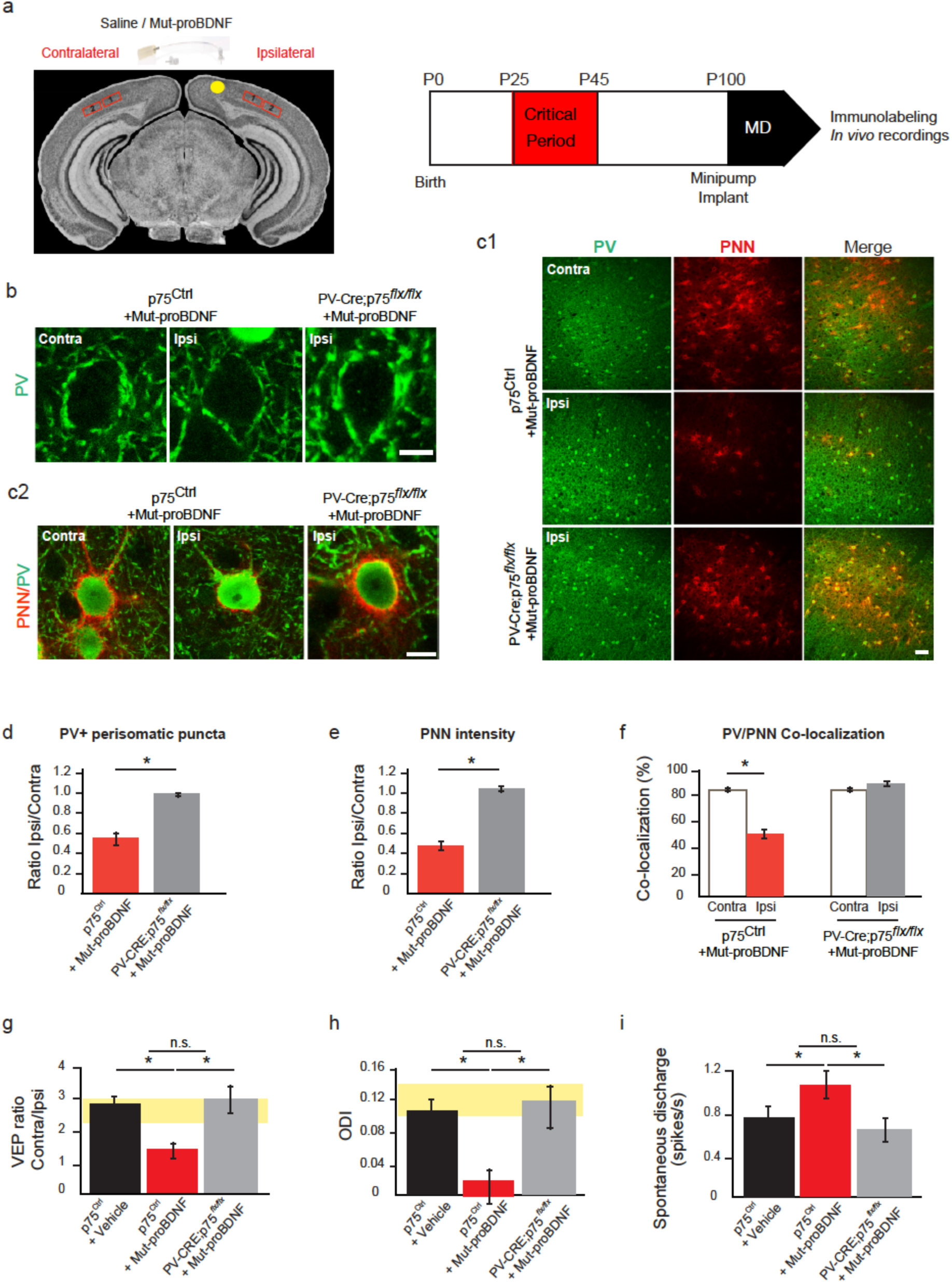
proBNDF-mediated p75NTR activation in cortical PV cells reduces their perisomatic boutons and restores ocular dominance plasticity in adult visual cortex *in vivo*. (a) Experimental approach. (b) The number of immunolabeled PV-positive puncta (green) surrounding NeuN-positive neuronal somata (red) is reduced in the binocular visual cortex ipsilateral to the minipump releasing mut-proBDNF (Ipsi) compared to the contralateral cortex (Contra) in the same animal. On the other hand, the number of PV-positive puncta per NeuN-positive profile in the ipsilateral cortex of *PV-CRE; p75^flx/flx^* mice is similar to that observed in the contralateral, untreated cortex. (c) Low (c1) and high (c2) magnification of PNN (red, WFA staining) enwrapping PV cells (green) show a dramatic reduction in both PNN density and intensity in the visual cortex infused with mut-proBFNF. This effects is abolished in *PV-CRE; p75^flx/flx^* mice. Scale bar, c1: 100µm; b, c2: 10µm. (d) Quantification of the mean number of PV-positive puncta per NeuN-positive profile in ipsilateral compared to contralateral cortex. Ipsi/Contra ratio is obtained for each animal, and then averaged between different animals. Mean Ipsi/Contra ratio is significantly reduced in Mut-proBDNF infused *p75^Ctrl^* but not in *PV-CRE; p75^flx/flx^* mice (t-test, p<0.001). (e) Mean PNN intensity around PV cells is significantly lower in the ipsilateral cortex of *p75^Ctrl^* but not *PV-CRE; p75^flx/flx^* mice infused with mut-proBDNF (t-test, p<0.001). (f) The percentage of PV cells colocalizing with PNN is significantly reduced in the cortex infused with mut-proBDNF (Ipsi) compared to the untreated cortex (Contra) in *p75^Ctrl^* (t-test, p=0.002) but not *PV-CRE; p75^flx/flx^* (t-test, p=0.192). n= 5 *p75^Ctrl^* mice; n=3 *PV-CRE; p75^flx/flx^* mice. (g) Contralateral to ipsilateral eye (C/I) VEP ratio mean values. The grey area denotes the C/I VEP ratio range in adult binocular animals. Three days of MD did not affect the C/I VEP ratio in adult mice, whereas it led to a significant decrease in the C/I VEP ratio of animals treated with mut-proBDNF. Mut-proBDNF effects was however abolished in *PV-CRE; p75^flx/flx^* mice (one-way ANOVA, *post-hoc* Holm-Sidak, p<0.001). *p75NTR^Ctrl^* + vehicle: n = 9, *p75NTR^Ctrl^* + mut-proBDNF: n=8, *PV-CRE; p75^flx/flx^* +mut-proBDNF: n=7 PV-CRE. (h) Histogram represents the average ODI ± SEM for each experimental group. The grey area defines the range of typical values for binocular adult animals. ODIs of *p75NTR^Ctrl^* mice infused with vehicle solution and *PV-CRE; p75^flx/flx^* mice infused with mut-proBDNF are not significantly different from those of undeprived animals, while ODIs in *p75^Ctrl^* mice treated with mut-proBDNF are significantly shifted towards the open eye (Kruskal-Wallis One Way ANOVA vs control, *post hoc* Dunn’s test, p<0.05). (i) Mean spontaneous discharge is significantly increased only in *p75^Ctrl^* mice treated with mut-proBDNF (Kruskal-Wallis One Way ANOVA, *post hoc* Dunn’s test, p<0.001). *p75NTR^Ctrl^* + vehicle: n = 9 mice, 174 cells; *p75NTR^Ctrl^* + mut-proBDNF: n=7 mice, 147 cells; *PV-CRE; p75^flx/flx^* +mut-proBDNF: n=6 mice, 125 cells.

To determine whether mut-proBDNF action was mediated through p75NTR activation in adult cortical PV cells, we infused the recombinant mutant protein in the visual cortex of conditional mutant mice (PV_Cre;p75NTR^lox/lox^). By introducing the RCE^EGFP^ allele to drive EGFP expression in presence of Cre, we showed that around 90% of PV cells co-expressed GFP (92 ± 1%; n = 4 mice), while virtually all GFP+ cells expressed PV in the adult (>P60) visual cortex of PV_Cre;p75NTR^lox/lox^ mice. Interestingly, PV_Cre;p75NTR^lox/lox^ adult mice did not show any significant difference in the number and intensity of PV+ puncta formed around pyramidal cells compared to their control littermates (p75NTR^Ctrl^) (perisomatic PV ring intensity: 84 ± 6 and 67 ± 9 a. u., number of perisomatic PV+ puncta: 7.9 ± 0.9 vs. 9.2 ± 0.2 for p75NTR^Ctrl^ vs. PV_Cre;p75NTR^lox/lox^, respectively; unpaired t-test, p > 0.1; n = 4 PV_Cre;p75NTR^lox/lox^ and n=3 p75NTR^Ctrl^ mice). In addition, visual cortex functional properties, analyzed by optical imaging, were not altered in adult PV_Cre;p75NTR^lox/lox^ mice compared to wild-type littermates (Supplementary Fig. 7). Cre expression occurs slowly and starts well after the second postnatal week in this mouse line^41^, thus it is possible that p75NTR knockout may occur too late to influence the development of PV cell connectivity. Another possibility is that p75NTR deletion might cause an acceleration of PV cell synapse maturation, which would have reached plateau by adulthood. Thus, since our analysis was performed in adult mice, we could not detect any difference between the two genotypes. Nonetheless, in contrast to what we observed following mut-proBDNF in control mice, mut-proBDNF infusion in mutant mice was unable to significantly alter perisomatic PV+ and VGAT+ puncta density (Fig. 7b, d; Supplementary Fig. 6), indicating that the effect of mut-proBDNF on perisomatic GABAergic boutons in adult mice was mediated by p75NTR expressed by PV cells. All together, these data suggest that activation of p75NTR onto PV cells mediated by pharmacological proBDNF treatment is able to destabilize PV cells connectivity in the adult brain.

### proBDNF-mediated p75NTR activation in PV cells promotes cortical plasticity in adult mice

Using ocular dominance plasticity in visual cortex as experimental model, recent studies showed that modulation of inhibition in adult brain can re-activate juvenile-like cortical plasticity mechanisms^13^. Since our data showed that treatment with mut-proBDNF could destabilize PV cell innervation in the adult brain *in vivo*, we asked whether this could in turn promote cortical plasticity. To answer this question, we first analyzed PNN expression pattern in visual cortex of mice infused with mut-proBDNF (Fig. 7a), since it has been shown that PNN normally enwrap mature PV cells to limit adult plasticity^43,44^. In p75NTR^Ctrl^ mice, mut-proBDNF infusion significantly reduced both the number of PV cells that were encircled by PNN, as revealed by WFA staining (Fig. 7c1, f; unpaired t-test, p = 0.002) and PNN immunofluorescence intensity around single PV+ cells (Fig. 7c2, e; ∼55% reduction in ipsi-vs contralateral cortex; unpaired t-test, p < 0.001). The effects of mut-proBDNF treatment on PNN were completely abolished in PV_Cre;p75NTR^lox/lox^ mice (Fig. 7c1-2, e, f). Importantly, PNN staining did not differ between untreated PV_Cre;p75NTR^lox/lox^ mice and control littermates (PNN intensity; 143 ± 6 and 148 ± 5 a. u., percentage of PV cells encircled by PNN: 87.6 ± 1.0 and 87.1 ± 0.9%, for p75NTR^Ctrl^ and PV_Cre;p75NTR^lox/lox^, respectively; unpaired t-test, p > 0.1, n = 4 PV_Cre;p75NTR^lox/lox^ and n = 3 p75NTR^Ctrl^ mice), suggesting that p75NTR activation by mut-proBDNF treatment was the critical step leading to PNN reduction.

To directly test whether mut-proBDNF could reopen a window of plasticity in adult visual cortex, we performed electrophysiological recordings in binocular regions of the primary visual cortex following a brief (3 days) monocular deprivation in adult mice. During the critical period for ocular dominance plasticity, the ratio of the amplitudes of visual evoked potentials (VEPs) evoked by eye stimulation shifts in favor of the non-deprived eye (ocular dominance shift). However no significant ocular dominance shift can be observed following three days of monocular deprivation at or after P100^43,45^. Consistently, we found that monocular deprivation did not affect the contralateral/ipsilateral (C/I) VEP ratio in vehicle-treated animals with respect to p75NTR^Ctrl^ non-deprived mice, while in p75NTR^Ctrl^ mice treated with mut-proBDNF we observed a marked ocular dominance shift in favor of the non-deprived eye, reflected by a significant decrease of the C/I VEP ratio (Fig. 5g; Supplementary Fig. 8a; One-way ANOVA, *post-hoc* Holm-Sidak, p < 0.001). p75NTR deletion in PV cells (PV_Cre;p75NTR^lox/lox^) completely prevented the ocular dominance shift induced by mut-proBDNF treatment (Fig. 7g, Supplementary Fig.8a).

To further confirm this data, we performed single-unit recordings. Ocular dominance of cortical neurons was assessed by quantitative evaluation of responsiveness to optimal visual stimulation of either eye and an ocular dominance index (ODI) was assigned to every single cell recorded. ODI of vehicle-infused, monocularly deprived, p75NTR^Ctrl^ mice displayed the typical bias towards the contralateral eye inputs as shown by non-deprived control mice, while mut-proBDNF infused, monocularly deprived, p75NTR^Ctrl^ mice showed a prominent ocular dominance shift in favor of the open eye (Fig. 7h, Supplementary Fig. 10b; Kruskal-Wallis One-way ANOVA vs control, *post hoc* Dunn’s test, p < 0.05), which was abolished in PV_Cre;p75NTR^lox/lox^ mice (Fig. 5h, Supplementary Fig. 8b). ODI values were significantly lower in mut-proBDNF-infused, monocularly deprived, p75NTR^Ctrl^ mice as compared with vehicle-infused, monocularly deprived, p75NTR^Ctrl^ mice and PV_Cre;p75NTR^lox/lox^ (Fig. 5h, Supplementary Fig. 8b; Kruskal-Wallis One Way ANOVA, *post hoc* Dunn’s test, p < 0.05). Consistently, the ODI cumulative distribution did not differ between no-monocularly deprived, and vehicle-treated, monocularly deprived, p75NTR^Ctrl^ mice (Supplementary Fig. 8c1; K-S test, p = 0.541), whereas both groups differed from p75NTR^Ctrl^ mice treated with mut-proBDNF (Supplementary Fig. 8c1; K-S test, p < 0.05). ODI distribution of mut-proBDNF-treated, monocularly-deprived PV_Cre;p75NTR^lox/lox^ was indistinguishable to that of non-deprived mice (Supplementary Fig. 8c2; K-S test, p = 0.633), whereas it was statistically different for monocularly-deprived p75NTR^Ctrl^ mice treated with mut-proBDNF (Supplementary Fig. 8c2; K-S test, p < 0.05), further proving that mut-proBDNF treatment was able to induce visual experience-dependent plasticity only in mice carrying intact p75NTR expression in PV cells.

Interestingly, we found that spontaneous discharge of visual cortical neurons was increased by mut-proBDNF treatment only in p75NTR^Ctrl^ mice (Fig. 7i; Kruskal-Wallis One Way ANOVA, *post hoc* Dunn’s test, p < 0.001), supporting the hypothesis that proBDNF-mediated p75NTR activation in PV cells reduces intracortical inhibition in adult mice.

In summary, these data demonstrate that p75NTR activation in cortical PV cells induces loss of PV cell connectivity and restoration of juvenile-like level of ocular dominance plasticity in adult mice.

## DISCUSSION

In this study we focused on the role of p75NTR in regulating interneuron synapse maturation during development and adult visual cortical plasticity. We had to first overcome the technical challenge of visualizing the presence of p75NTR in PV neurons during development and in the adult visual cortex, the focus of this study. Using two cutting-edge experimental approaches to detect very low levels of RNA and protein, both in the developing and adult cortex, we were able to specifically detect p75NTR in not just PV somata but also in presynaptic terminals. Next, we showed that p75NTR expression levels and activation can modulate the formation of PV cell connectivity during development in organotypic slice cultures and *in vivo.* Finally, we proved that pharmacological activation of p75NTR in PV cells reduces PV cell connectivity and allows juvenile-like plasticity, in adult visual cortex.

During development, p75NTR is slowly downregulated after the third postnatal week, while at the same time PV cells develop complex, highly branched axonal arbors that contact an increasingly higher number of potential postsynaptic targets^15^. Our results show that PV cell-specific p75NTR gene loss accelerates, whereas p75NTR activation hinders the development of complex perisomatic innervation fields. Therefore, p75NTR acts as a negative signal constraining the formation of PV cell connectivity. Many studies have addressed the molecular signals promoting the development of PV cell innervations^16–18,20^; on the other hand the involvement of factors constraining PV cell innervation field is less well understood. The polysialic acid motif PSA was previously shown to hinder PV cell synapse formation before eye opening^10^. PSA is a general modulator of cell interactions and, as such, it likely acts as a permissive signal to allow optimal interactions between presynaptic PV axons and postsynaptic cells. On the other hand, here, our data show that p75NTR expression levels specifically in PV cells negatively regulates the extent of its innervations field. Therefore, it is possible that p75NTR expression level may act as an instructive signal for PV cell innervation refinement. In fact, using PLA, we found that a population of PV cell boutons colocalizes with p75NTR. Locally, p75NTR activation may inhibit the formation of PV cell innervation by promoting growth cone collapse, via activation of RhoA^46,47^ and/or inactivation of Rac signaling, which leads to destabilization of actin filaments and collapse of neurite outgrowth^48^. Further, it has been suggested that p75NTR activation may sensitizes neurons to other inhibitory, growth cone collapsing cues such as Nogo^49,50^, ephrins and semaphorins^46,51^. It will be interesting to study whether and how these inhibitory cues interact with p75NTR signaling to modulate the maturation of PV cell innervation. In addition to locally regulating cytoskeletal dynamics, p75NTR activation can cause changes in gene transcription, leading to modulation in expression of proteins modifying PV cell synaptic inputs and/or excitability, which would in turn regulate PV cell axon growth^52^.

There are open questions regarding mechanisms regulating p75NTR downregulation during development. A recent study suggests that p75NTR expression level is negatively regulated by visual experience *in vivo*^27^. In fact, Bracken and Turrigiano^27^ have shown that, in visual cortex, p75NTR mRNA levels strongly decrease after eye opening (around the second postnatal week), and p75NTR, but not TrkB, mRNA levels are upregulated by prolonged dark rearing. It is conceivable that activity levels in individual PV cells regulate their p75NTR expression, which in turn determines to what extent they respond to local changes in molecular p75NTR regulators. In addition, p75NTR expression is regulated by early growth response (Egr) factors 1 and 3, which are inducible transcriptional regulators modulating gene expression in response to a variety of extracellular stimuli influencing cellular growth, differentiation and response to injury^53^, suggesting a potentially highly dynamic, and cell context-dependent mechanism for regulation of p75NTR expression during development or following injury. In accordance with this hypothesis, it has been shown that p75NTR is upregulated by pathological events, including cerebral ischemia^54^ and seizures^55,56^. One implication of our findings is that pathology-induced upregulation of p75NTR levels occurring during early brain development impair the maturation of PV cell circuits, which may in turn affect the expression and/or timing of critical period plasticity^10,11^, thus contributing to long-term cognitive and behavioral impairments.

In the adult, the brain’s intrinsic potential for plasticity is actively dampened, by increase in intracortical inhibition and the simultaneous expression of brake-like factors, which limit experience-dependent circuit rewiring beyond a critical period. Interestingly, many of these plasticity breaks converge onto PV cell function^14,43,57^. Our results demonstrate that reducing PV cell connectivity is sufficient to promote juvenile-like levels of ocular dominance plasticity in the adult cortex. p75NTR activation may directly affect the stability of PV cell axonal branches and synapses, by affecting local cytoskeletal dynamics^47,48^. In addition, p75NTR activation can affect the synthesis of specific proteins, including those required for PNN condensation around PV-positive cells^58^. Intact PNNs structurally limit synaptic rearrangements of inputs onto PV cells, which in turn regulate their excitability and synaptic release. Consistently, reduction of PV cell excitability leads to a reduction of their innervation fields even after reaching maturity^52^. Further, PNN disruption may prevent the persistent uptake of the homeoprotein Otx2 into PV cells, which is required by the PV cells for the maintenance of an adult phenotype^11,14^.

In our study, we did not find significant differences in PV cell perisomatic connectivity, PNN intensity and visual cortical properties in the visual cortex of adult littermate p75NTR^lox/lox^ vs PV_Cre;p75NTR^lox/lox^ mice. One possibility is that Cre-mediated p75NTR knockout may occur too late to affect the maturation of the functional properties of the visual cortex, which reaches a plateau well before the onset of adolescence. Since p75NTR expression differs among brain regions at the different ages^23,24^, it would be interesting to investigate whether PV_Cre;p75NTR^lox/lox^ cKO mice show altered cognitive functions implicating regions which mature later, such as the prefrontal cortex and frontolimbic circuitry^23^.

The role of neurotrophins and their precursor forms in p75NTR-mediated signaling has been the subject of several debates. Numerous studies have shown that proNGF and proBDNF can promote cell death by interacting with a receptor complex consisting of p75NTR and sortilin (sortilin-related VPS10 domain-containing receptor)^59,60^ and that the extracellular conversion from proBDNF into BDNF promotes LTD in the hippocampus, by activating p75NTR^35,38^. In addition, while it was well accepted that the pro-domain plays a role in the folding, stability and intracellular trafficking of BDNF^61^, recent data has started to highlight the possibility that the BDNF prodomain *per se* may have diverse biological functions. Indeed, several recent reports indicated that the BDNF pro-domain is endogenously present and has biological effects. First, Dieni et al.^32^ reported that BDNF and its pro-peptide both stained large dense core vesicles in excitatory presynaptic terminals of the adult mouse hippocampus. Second, Mizui et al.^62^ showed that the BDNF pro-peptide facilitates LTD in the hippocampus. Third, Anastasia et al.^22^ showed that the prodomain is detected at high levels in the hippocampus *in vivo*, in particular after the first postnatal month, and that its secretion is activity-dependent in hippocampal neuronal cultures. Based on the relative expression of proBDNF, mBDNF and the BDNF prodomain during development and in the adult brain^22,31–33^, it has been hypothesized that secreted proBDNF may play a role during early development while the secreted prodomain may have biological effects in the adolescent and adult brain^30^. Consistent with his hypothesis, our data show that modulating endogenous mBDNF levels by acting on tPA activity before the third postnatal week affects the development of PV cell innervation and that this depends on p75NTR expression by PV cells. It remains to be established whether BDNF prodomain plays a role in the maintenance and plasticity of PV cell connectivity in developing and adult brains.

A common single-nucleotide polymorphism (SNP) in the human *BDNF* gene results in a Val66Met substitution in the BDNF prodomain region which is associated with impairments in specific forms of learning and memory and with enhanced risk of developing depression and anxiety disorders in humans and mice ^63–65^. In the light of these observations, it is interesting to note that Met66, but not Val66, prodomain is sufficient to induce neurite retraction in cultured hippocampal neurons in presence of both SorCS2 (sortilin-related Vps10p-domain sorting receptor 2) and p75NTR^22^ and to trigger mature mushroom spines elimination in the ventral hippocampus *in vivo*^23^. Since at least a subset of PV cells express p75NTR even in adulthood, it will be interesting to investigate whether the presence of the Met66 variant alters the formation and/or plasticity of PV cell innervation, thereby contributing to the endophenotypes related to neuropsychiatric disorders associated with the Val66Met polymorphism in humans.

## Acknowledgments

We thank Drs. Graçiela Pineyro (CHU Ste. Justine, Montreal), Elsa Rossignol (CHU Ste. Justine, Montreal), Edward Ruthazer (Montreal Neurological Institute, Montreal), Dr. Keith Murai (CRN, McGill University, Montreal) and Dr. Agustin Anastasia (INIMEC-CONICET –Universidad Nacional de Cordoba, Argentina) for their insightful suggestions, Dr. Louis Reichardt (USF) for providing reagents, Dr. Vesa Kartineen (UMichigan) for the p75NTR^lox^ mouse, Dr. Phil Barker (UBC, Vancouver) for the anti-p75NTR antibody and JF Cloutier (MNI, Montreal) for the p75NTR KO mice. We would like to thank Josianne Nuñes Carriço and Menna Murrah for their technical assistance as well as the Comité Institutionnel de Bonne Pratiques Animales en Recherche (CIBPAR) and all the personnel of the animal facility of the Research Center of CHU Sainte-Justine (Université de Montreal) for their instrumental technical support. This work was supported by the Canadian Institutes of Health Research (G.DC), Canada Foundation for Innovation (G.DC), Canada Research Chair Program (G.DC). E.B. and M.J-L are supported by FRQS fellowship. B.C. is supported by CHU Sainte-Justine Foundation.

## Materials and Methods

### Mice

Organotypic cortical cultures were prepared from C57Bl6 (Jackson Labs) or p75NTR^lox/lox^ mice (^28^, kindly provided by Dr. Vesa Kaartinen, University of Michigan). In this mouse, exons 4-6 of p75NTR, which encode the transmembrane and all cytoplasmic domains, are flanked by two loxP sites. PV_Cre; p75NTR^lox/lox^ mice were generated by crossing p75NTR^flx^ with PV_CRE mice (B6.129P2-Pvalb^tm1(cre)Arbr^/J; Jackson Laboratory). Cell-specificity of Cre-mediated recombination was analyzed by breeding PV_Cre^41^ with RCE^EGFP^ mice (*Gt(ROSA)26Sor^tm1.1(CAG-EGFP)Fsh^*/J; Jackson laboratory). This latter line carries a loxP-flanked STOP cassette upstream of the EGFP gene. Removal of the loxP-flanked STOP cassette by Cre-mediated recombination drives EGFP reporter expression. p75NTR^lox/lox^ and p75NTR^+/+^ mice were analyzed separately in all performed experiments; however, as we did not find any difference between these two genotypes (t-test or Mann Whitney test, p > 0.1), we pooled them together and indicated them as p75NTR^Ctrl^.

### Cortical organotypic culture and biolistic transfection

Slice culture preparation was performed as in^15,17^ using mice pups of either sex. Briefly, postnatal day 3 (P3) to P5 mice were decapitated, and brains were rapidly removed and immersed in culture medium (containing DMEM, 20% horse serum, 1 mM glutamine, 13 mM glucose, 1 mM CaCl_2_, 2 mM MgSO_4_, 0.5 µm/ml insulin, 30 mM HEPES, 5 mM NaHCO_3_, and 0.001% ascorbic acid). Coronal brain slices, 400 µm thick, were cut with a chopper (Stoelting, Wood Dale, IL). Slices were then placed on transparent Millicell membrane inserts (Millipore, Bedford, MA), usually 2-4 slices/insert, in 30 mm Petri dishes containing 750 µl of culture medium. Finally, they were incubated in a humidified incubator at 34°C with a 5% CO_2_-enriched atmosphere, and the medium was changed three times per week. All procedures were performed under sterile conditions. Biolistic transfection was performed as described in^17^. Constructs were incorporated into “bullets” that were made using 1.6 µm gold particles (Bio-Rad) coated with 25-30 µg of the each of the plasmids of interest. When a gold particle coated with multiple constructs enters the neuron, all the constructs are co-expressed within the same cell since they are driven by the same P_G67_ promoter. P_G67__GFP was originally generated by subcloning of a 10 kb region of *Gad1* gene promoter by gap repair in front of the GFP coding region in pEGFP (Clontech)^15^. Bullets were used to biolistically transfect organotypic slices using a gene gun (Bio-Rad, Hercules, CA) at high pressure (180Ψ), and the transfected slices were incubated for 6-8 days, under the same conditions as described above, before imaging. To label control PV cells, slices were transfected with P_G67__GFP bullets, while for the p75NTR^-/-^ PV cells were transfected with both P_G67__GFP and P_G67__Cre.

wt-proBDNF and mut-proBDNF (10 ng/ml, Alomone Labs) were respectively added with the culture medium during the specific time window indicated in the results section. To block p75NTR, disrupt tPA-induced endogenous proBDNF cleavage or overexpress tPA, REX antibody (50 μg/ml, Dr. Louis Reichardt, USF), PPACK peptide (50 μM, Molecular Innovations) and active tPA recombinant protein (0.6 μg/ml, Molecular Innovations) were respectively added within the culture medium. Every experimental data was repeated at least twice, using culture batches prepared in different days.

### Analysis of PV cell innervation

Previous studies have shown that the basic features of maturation of perisomatic innervation by PV-positive basket interneurons (referred as PV cells) onto pyramidal cells are retained in cortical organotypic cultures. In organotypic cultures, PV cells start out with very sparse and simple axons, which develop into complex, highly branched arbors in the subsequent 4 weeks with a time course similar to that observed *in vivo*^15^. We have previously shown that the vast majority of GFP-labeled boutons in our experimental condition most likely represent presynaptic terminals^10,16,17^.

For each experimental group, we took care to acquire an equal number of PV cells localized in layer 2/3 and 5/6. In average, we acquired only one PV cell from each successfully transfected organotypic culture. Confocal images of the PV cell axon arbors were taken in the first 150 μm from the PV cell soma using a 63X glycerol objective (NA 1.3, Leica) and a Leica SPE. Analysis of PV basket cell perisomatic innervation was performed as described in ^17^. Pyramidal cell somata were identified by NeuN immunofluorescence and the axon of PV cells were traced in 3D. Only innervated NeuN-positive cells were included in this analysis. The following parameters were analyzed for each PV cell: a) perisomatic bouton density, b) axonal terminal branching around innervated somata and c) percentage of pyramidal somata innervated by basket cell. In our 3D Sholl analysis, sholl spheres with a 1µm increment from the center of a pyramidal soma were used to quantify PV cell axon terminal branch complexity and bouton density around the pyramidal cell soma. Axon branch complexity around a single pyramidal cell soma was quantified by the average number of intersections between PV cell axons and the sholl sphere in the first 9 µm from the center of the pyramidal cell soma. We choose 9 µm as the limiting radius for a sholl sphere because it approximates the average pyramidal cell soma diameter measured from pyramidal neurons immunostained with NeuN antibody. Between 10 and 15 pyramidal neurons were analyzed for each basket cell. To quantify the fraction of pyramidal cell somata potentially innervated by a PV cell axon, we divided the number of NeuN-positive neurons contacted by at least one GFP positive-bouton by the total number of NeuN-positive cells, in a confocal stack (at least 2 stacks per PV cell). We measured NeuN-positive cell density and found it to be invariant with respect to the different manipulations. All data were first averaged per PV cell, thus statistical analysis was done using the number of PV cells as “n”.

### Western Blots

Membranes were probed with anti-mBDNF (1:200; Santa Cruz, N20: sc-546) and antiglyceraldehyde-3-phosphate dehydrogenase, 1:8000 (GAPDH, mouse monoclonal IgG; Cat. no. AM4300; Applied Biosystems, Streetsville, Ontario, Canada). Each sample corresponded to 6 organotypic cultures pooled together. In addition, Ctrl samples were collected for each mouse used for organotypic cultures. All samples used for western blot analysis of a specific protein were run on the same gel. Membranes were exposed to Bioflex MSI autoradiography / X-ray film for different time intervals, and only the films that showed easily identifiable, but not saturated, bands for every sample were used for quantification, using imageJ software (Wayne Rasband, National Institutes of Health, USA, http://imagej.nih.gov/ij). Background mean grey value was subtracted and the values were normalized on GAPDH mean grey value. The average of normalized mean grey value of control experiments was calculated and assigned a value of 1. The normalized values of the PPACK and tPA treatments were then expressed as the relative of the control experiments. Specificity of the anti-BDNF antibody was verified using brain lysates from CaMKII_Cre;BDNF^lox/lox^ and their BDNF^lox/lox^ adult littermates (Supplementary Fig. 3).

In addition, we tested the following anti-proBDNF antibodies: chicken anti-proBDNF (Millipore, AB9042), rabbit-anti-proBDNF (Alomone Labs, ANT-006) and guinea-pig-anti-proBDNF (Alomone Lab, AGP-032). However, in our hands, we could still detect the proBDNF band in lysates from CaMKII_Cre; BDNF^lox/lox^ mice, therefore we could not confirm their specificity and did not use them further in our studies.

### Proximity Ligation Assays (PLA)

Mice were anesthetized and transcardially perfused with ACSF (Artificial CerebroSpinal Fluid). After extraction, brain were incubated at 4°C overnight in 4% paraformaldehyde. Sagittal sections, 60 µm thick, were blocked with 10 % horse serum and permeabilized with 0.2% Triton X-100 (v/v). Experiments were then performed according to the manufacturer’s instructions (Duolink^®^ & PLA^®^ Technology, Olink-Bioscience, Uppsala, Sweden).

Briefly, sections were incubated with goat-anti p75NTR antibody (R&D Systems, Cat#AF1157) at 4°C for 24-36 hours. PLA probes anti-goat plus and minus, which are secondary antibodies conjugated with oligonucleotides, were added and incubated for 1 h at 37°C. Amplification template oligonucleotides were hybridized to pairs of PLA and circularized by ligation. The hence formed DNA circle was then amplified using rolling circle amplification and detection of the amplicons was carried out using the 624 Duolink *in situ* detection kits, resulting in red fluorescence signals. Sections were mounted and were analyzed under a 40X oil immersion objective using a confocal microscope (Zeiss LSM 780 or Leica TCS SP8 X). Distinct bright spots contained within an area of the section designated by the experimenter were counted using an ImageJ macro. Briefly, we determined a pre-sized zone of interest (ROI) and then performed segmentation by thresholding in order to generate binary images from each selection. The number of individual points was quantified using the granulometry algorithm of ImageJ. Each experiment was repeated 3 times.

Specificity of anti-p75NTR antibodies was tested by performing immunofluorescence staining in an adult p75NTR KO mouse and its wild-type littermates, kindly provided by Dr. JF Cloutier (data not shown).

### Fluorescent multiplex RNAscope

To prepare tissue for *in situ* hybridizations (ISH), mice were anesthetized and perfused with saline (0.9% NaCl) followed by 4% paraformaldehyde/phosphate buffer, pH 7.4. Brains were dissected and post-fixed in 4% PFA for 24 hr at 4°C, cryoprotected first in 15% and then in 30% sucrose in PBS and embedded in OCT. Brain sections (15 µm) were cut using a cryostat (Leica) and mounted on superfrost plus gold glass slides (Fisher Scientific #22-035-813). Slides were subsequently stored at -80°C. Probes for Mm-Ngfr (494261), and Pvalb (421931-C2) as well as all other reagents for *in situ* hybridization, were purchased from Advanced Cell Diagnostics (ACD, Newark, CA). The tissue pretreatment, hybridization, amplification, and detection were performed according to User Manual for Fixed Frozen Tissue (ACD). During RNAscope hybridization, positive probes, negative probes and PV/p75 probes were processed simultaneously. Briefly, the slides were removed from -80C and rinsed with 1X PBS to remove OCT. After they were submerged into 1X Target retrieval solution for 5 min at 100°C, and then rinsed in distilled water followed by 100% EtOH dip to remove access water. Protease III was added to each section and incubated for 30 min at 40°C followed by washing in distilled water. For detection, probes were added to each section and incubated for 2 hr at 40°C. Unbound probes were subsequently washed away by rinsing slides in 1X wash buffer. AMP reagents were added to each section and incubated for as per manufacturer’s instructions, and washed in wash buffer for 2 min. Sections were stained with DAPI for 30 s, and then mounted with Prolong Gold Antifade Mountant. This experiment was performed using tissue from three different mice.

### Immunostaining analysis

Cortical organotypic cultures were fixed, freeze-thawed and immunostained as previously described^15^. Mice were perfused with saline followed by 4% paraformaldehyde in phosphate buffer (pH 7.4). Brains were then removed and post-fixed overnight at 4°C in the same fixative solution, cryoprotected in 30% sucrose in PBS for 1 to 2 days, then frozen in Tissue Tek. 40µm thick brain slices were obtained using a cryostat (Leica). The following antibodies were used: NeuN (mouse monoclonal, 1:400, Millipore; Cat#MAB377), PV (mouse monoclonal, 1:5000, Swant, Cat#235), PV (Rabbit polyclonal, 1:5000, Swant, Cat# PV25), VGAT (Rabbit polyclonal, 1:400, Synaptic Systems, Cat#131003), gephyrin (mouse monoclonal, 1:500, Synaptic Systems, Cat#147 021) followed by the appropriate Alexa555-conjugated or Alexa633-conjugated IgG (Molecular Probes, 1: 400).

To label PNN, brain slices were incubated in a solution of biotin-conjugated lectin *Wisteria floribunda* (WFA) (10 µg/ml; Sigma-Aldrich) followed by Alexa 568-conjugated extravidin (1:500 in PBS; Sigma-Aldrich). Tissue was mounted in Vectashield mounting medium (Vector) before imaging.

### Immunolabeling imaging and analysis

Mice were anesthetized and perfused with saline (0.9% NaCl) followed by 4% paraformaldehyde/phosphate buffer, pH 7.4, then the brain was extracted and cryoprotected in 30% sucrose/PBS, and frozen in Tissue Tek. For PV, vGAT and PNN analysis on minipump-implanted brains, sections were processed in parallel and images were all acquired the same day using identical confocal parameters. Confocal images (Leica, SPE or Leica SP8) were acquired using either a 20x water immersion objective (NA 0.7; Leica) or a 63x glycerol objective (NA 1.3; Leica). For each animal, we acquired two confocal stacks in layer 5 in both hemispheres (infused, Ipsi vs non-infused, Contra). Data were obtained from 3 to 4 brain sections per animal. Z-stacks were acquired with a 1 μm step, exported as TIFF files, and analyzed using ImageJ software. PV, vGAT or PNN perisomatic rings (between 7 to 10 in each stack) were outlined and the mean gray values were measured, after background subtraction.

For PV/gephyrin puncta analysis, confocal images (Leica SP8) were acquired using a 63x glycerol objective (NA 1.3; Leica). For each animal, we acquired one confocal stack with a 0.3 μm step in cortical layer 5 from 3 to 4 brain sections per animal. Stacks were exported as TIFF files and analyzed using ImageJ software. All analysis was done by operators blind to the mouse genotype or to the specific treatment.

### Minipump implant and Monocular Deprivation (MD)

Adult (>P100) mice were implanted with osmotic mini-pump under isoflurane anesthesia. Minipumps (model 1007D; flow rate 0.5 µl/h; Alzet) were filled with mut-proBDNF (1 µg/ml in filtered PBS, Alomone Laboratories) or vehicle solution and connected to a cannula (gauge 30) implanted directly in the primary visual cortex (2.5 mm lateral to the midline, 2.5 mm anterior to lambda).

For electrophysiological analysis, a group of animals were monocularly deprived through eyelid suturing two days after the implant of the minipump, and then recorded 3 days after. Subjects with even minimal spontaneous re-opening were excluded from the study. For perisomatic GABAergic bouton density and PNN studies, a second group of animals was perfused 5 days after minipump implant.

### In Vivo Electrophysiology

After 3 days of MD, animals were sedated with isoflurane and anesthetized with urethane (i.p. injection; 1.5 g/kg; 20% solution in saline; Sigma, St. Louis, MO, USA), then placed in a stereotaxic frame. Body temperature was maintained at 37°C. A hole was drilled in the skull, corresponding to the binocular portion of the primary visual cortex (binocular area Oc1B), contralateral to the deprived eye. Dexamethasone (2 mg/kg) was administered subcutaneously to reduce secretions and edema and saline was periodically infused to prevent dehydration. Eyes were covered with a thin layer of silicone oil to avoid corneal opacities. Recordings were made using silicon microprobes (16 channels, NeuroNexus Technologies a2×2-tet-3mm-150-121) inserted into the cortex 3.0-3.2 mm from the lambda point. Signals were acquired using Cheetah 5 (Neuralynx) and analyzed with custom software in Matlab (MathWorks).

#### Visual Stimulation

Stimuli were generated in Matlab using the Psychophysics Toolbox extensions and displayed with gamma correction on a monitor (Sony Trinitron G500, 60 Hz refresh rate, 32 cd/m2 mean luminance) placed 20 cm from the mouse, subtending 60-75° of visual space.

#### Visual evoked potentials (VEPs)

VEP were recorded as described in^66^. We measured contralateral to ipsilateral ratio of VEP amplitude to measure ocular dominance plasticity. Extracellular signal was filtered from 1 to 275 Hz. VEPs in response to square wave patterns with a spatial frequency of 0.06 cpd and abrupt phase inversion (1 Hz temporal period), were evaluated in the time domain by measuring the P1 peak-to-baseline amplitude and latency. Computer controlled mechanical shutters were used to collect data from each eye.

#### Single-Units

For single-unit recording extracellular signal was filtered from 0.6 to 6 kHz. Sampling rate: 33 kHz. Spiking events were detected on-line by voltage threshold crossing and waveforms of 1 ms were acquired around the time of threshold crossing. To improve isolation of units, recordings from groups of four neighboring sites (tetrode) were linked, so that each spike was composed by 4 waveforms. Then waveforms were processed using the OffLine Sorter software (Plexon). Drifting sinusoidal gratings were used as visual stimuli (1.5 s duration, temporal frequency of 2 Hz, 12 directions, 6 spatial frequency: 0.01, 0.02, 0.04, 0.08, 0.16, 0.32 cpd). Stimulation was repeated five times per eye, with stimulus conditions randomly interleaved, and two gray blank conditions (mean luminance) were included in all stimulus sets to estimate the spontaneous firing rate.

The average spontaneous rate for each unit was calculated by averaging the rate over all blank condition presentations. Responses at each orientation and spatial frequency were calculated by averaging the spike rate during the 1.5 s stimulus presentations and subtracting the spontaneous rate. The preferred stimulus was determined finding the combination of spatial frequency and orientation that maximize the response, independently for each eye. Ocular Dominance Index (ODI) was calculated as follows: ODI = (respContra-respIpsi)/(respContra+respIpsi), where ‘resp’ is the response evoked by the preferred stimulus, ‘Contra’ and ‘Ipsi’ are respectively: contralateral and ipsilateral eye. Experiments were done by operators blind to the genotype.

### In Vivo Optical Imaging

Optical imaging experiments were performed as in ^67^. Briefly, mice were anesthetized with urethane (1.25 g/kg, i.p.). Core body temperature was maintained at 37 °C using a feedback controlled heating pad (Harvard Apparatus, Saint-Laurent, Québec) and electrocardiogram (FHC, Bowdoin, ME, USA) was continuously monitored with sub-dermal electrodes. The visual cortex was imaged through the skull: an imaging chamber was placed over both hemispheres, glued on the skull, filled with agarose (1%) and sealed with a coverslip.

#### Stimulation

Visual stimulation was provided using VPixx and presented by an LCD projector on a screen placed at a distance of 20 cm in front of the mouse eyes (subtending 150 × 135° of visual angle). To assess visuotopy and characterize maps and connectivity in V1, we used a continuous stimulation paradigm, where 2° thick light bars were periodically shifted horizontally (to obtain elevation maps) or vertically (to obtain azimuth maps) over a dark background at a frequency of 0.15 Hz. These relative retinotopic maps were used to assess several structural and functional parameters within V1. To examine the functional properties of V1 neurons, episodic full-field sine wave grating stimuli (270°) were presented during 2 s and spaced by a blank presentation lasting 18 s intervals (mean luminance 75 cd/m^2^). The amplitude of the hemodynamic responses was measured as a function of contrast and spatial frequency selectivity. Five contrasts (6%, 12%, 25%, 50% and 90%) and seven spatial frequencies (0.01, 0.025, 0.05, 0.12, 0.24, 0.32 and 0.48 cycle per degree (cpd)) were used to determine contrast sensitivity and spatial frequency selectivity, respectively.

#### Image acquisition

The cortex was illuminated at 545 nm to adjust the focus of the camera and at 630 nm to record the intrinsic signals. Optical images were recorded using a 12-bit CCD camera (1M60, Dalsa, Colorado Springs, USA) driven by the Imager 3001 system (Optical Imaging Inc.©) and fitted with a macroscopic lens (Nikon, AF Micro Nikon, 60 mm, 1:2:8D). Frames of 512 × 512 pixels were acquired at a rate of 4 Hz, giving a spatial resolution of 28 µm/pixel. The acquisition was sustained for 10 min during the continuous stimulation paradigm. During episodic stimulation, frames were acquired for 20 s for every contrast and spatial frequency tested. An average of 10 repetitions was used to obtain a good signal to noise ratio.

#### Data analysis

OIS data were analyzed with MATLAB (MathWorks, Nattick, MA). For each pixel of the cortex, a Fourier transform was applied on temporal signals collected during continuous stimulation. Fourier phase and amplitude were generated for each frequency and used to map the retinotopy and realize quantification. The amplitude of neuronal activity was used to generate the “neuronal activation” map. In parallel, the phase at the stimulus frequency was related to the delay to activate the receptive field and was associated to the relative retinotopic position. The “retinotopic” map was obtained by multiplying the amplitude and phase maps. Regions of interest (ROI) located in the occipital cortex were manually delineated in the activation maps for each hemisphere. The area of V1 was calculated from the ROI borders. The shape of the ROI was fitted to an ellipse with MATLAB and the ratio of length of the two main axes of the ellipse determined (height/width) was calculated to measure the “ovality index”. The ratio of the number of the phases detected in the retinotopic maps over 2π (i.e. the range of the phases displayed) was used to estimate the “apparent visual field”, i.e. the proportion of the activated visual field represented in V1. The difference between the phase of each pixel and its surrounding pixels was calculated on the phase map to evaluate the “scatter index”. Fourier amplitude at the stimulus frequency and second harmonic was used to evaluate the population receptive field (pRF) size of the underlying neurons (neurons within a ROI respond to a range of visual field locations and the region of the visual space that stimulates this local neuronal activity is called pRF).

The hemodynamic responses obtained during episodic stimulation were used for the functional analysis of the neuron features. The contrast and spatial frequency tuning curves for each pixel of V1 were established from the amplitude of the negative peak of the hemodynamic response. The spatial frequency producing the strongest hemodynamic response was calculated for each pixel. For each animal, the results of each trial were pooled and an asymmetric Gaussian curve was fitted on the normalized values. Curves that did not meet the p < 0.05 and r-square ≥ 0.700 were not used. The optimal spatial frequency was defined as the spatial frequency producing the strongest response. The visual acuity was measured using a linear fit. The curves of amplitude as a function of the contrast were fitted with a Naka-Rushton function to determine the contrast evoking 50% of the maximum response.

### Statistics

Data were expressed as mean ± SEM unless otherwise specified in the legends. Normality tests were performed for all data analyzed. Differences between two groups were assessed with the Student’s unpaired *t*-test for normally distributed data or with the Mann Whitney Rank Sum test for not-normally distributed data. Differences between multiple groups were assessed with one-way ANOVA, and the specific *post hoc* tests used are reported in the legends. Statistical analysis was performed using Prism 7.0 (GraphPad Software). No animal was excluded from the analysis.

### Data availability

Detailed statistics and data that support the findings of this study are available from the corresponding authors on request.

## References

1. Cardin, J. A. et al. Driving fast-spiking cells induces gamma rhythm and controls sensory responses. Nature 459, 663–667 (2009).

2. Sohal, V. S., Zhang, F., Yizhar, O. & Deisseroth, K. Parvalbumin neurons and gamma rhythms enhance cortical circuit performance. Nature 459, 698–702 (2009).

3. Takada, N. et al. A developmental cell-type switch in cortical interneurons leads to a selective defect in cortical oscillations. Nat. Commun. 5, 5333 (2014).

4. Fries, P., Reynolds, J. H., Rorie, A. E. & Desimone, R. Modulation of oscillatory neuronal synchronization by selective visual attention. Science 291, 1560–1563 (2001).

5. Fries, P. Neuronal gamma-band synchronization as a fundamental process in cortical computation. Annu. Rev. Neurosci. 32, 209–224 (2009).

6. Howard, M. W. et al. Gamma oscillations correlate with working memory load in humans. Cereb. Cortex N. Y. N 1991 13, 1369–1374 (2003).

7. Cho, R. Y., Konecky, R. O. & Carter, C. S. Impairments in frontal cortical gamma synchrony and cognitive control in schizophrenia. Proc. Natl. Acad. Sci. U. S. A. 103, 19878–19883 (2006).

8. Fagiolini, M. & Hensch, T. K. Inhibitory threshold for critical-period activation in primary visual cortex. Nature 404, 183–186 (2000).

9. Fagiolini, M. et al. Specific GABAA circuits for visual cortical plasticity. Science 303, 1681–1683 (2004).

10. Di Cristo, G. et al. Activity-dependent PSA expression regulates inhibitory maturation and onset of critical period plasticity. Nat. Neurosci. 10, 1569–1577 (2007).

11. Sugiyama, S. et al. Experience-dependent transfer of Otx2 homeoprotein into the visual cortex activates postnatal plasticity. Cell 134, 508–520 (2008).

12. Kobayashi, Y., Ye, Z. & Hensch, T. K. Clock genes control cortical critical period timing. Neuron 86, 264–275 (2015).

13. Harauzov, A. et al. Reducing intracortical inhibition in the adult visual cortex promotes ocular dominance plasticity. J. Neurosci. Off. J. Soc. Neurosci. 30, 361–371 (2010).

14. Beurdeley, M. et al. Otx2 binding to perineuronal nets persistently regulates plasticity in the mature visual cortex. J. Neurosci. Off. J. Soc. Neurosci. 32, 9429–9437 (2012).

15. Chattopadhyaya, B. et al. Experience and activity-dependent maturation of perisomatic GABAergic innervation in primary visual cortex during a postnatal critical period. J. Neurosci. Off. J. Soc. Neurosci. 24, 9598–9611 (2004).

16. Chattopadhyaya, B. et al. GAD67-mediated GABA synthesis and signaling regulate inhibitory synaptic innervation in the visual cortex. Neuron 54, 889–903 (2007).

17. Chattopadhyaya, B., Baho, E., Huang, Z. J., Schachner, M. & Di Cristo, G. Neural cell adhesion molecule-mediated Fyn activation promotes GABAergic synapse maturation in postnatal mouse cortex. J. Neurosci. Off. J. Soc. Neurosci. 33, 5957–5968 (2013).

18. Del Pino, I. et al. Erbb4 deletion from fast-spiking interneurons causes schizophrenia-like phenotypes. Neuron 79, 1152–1168 (2013).

19. Huang, Z. J. et al. BDNF regulates the maturation of inhibition and the critical period of plasticity in mouse visual cortex. Cell 98, 739–755 (1999).

20. Kohara, K. et al. A local reduction in cortical GABAergic synapses after a loss of endogenous brain-derived neurotrophic factor, as revealed by single-cell gene knock-out method. J. Neurosci. Off. J. Soc. Neurosci. 27, 7234–7244 (2007).

21. Lin, Z. et al. Structural basis of death domain signaling in the p75 neurotrophin receptor. eLife 4, e11692 (2015).

22. Anastasia, A. et al. Val66Met polymorphism of BDNF alters prodomain structure to induce neuronal growth cone retraction. Nat. Commun. 4, 2490 (2013).

23. Giza, J. I. et al. The BDNF Val66Met Prodomain Disassembles Dendritic Spines Altering Fear Extinction Circuitry and Behavior. Neuron 99, 163-178.e6 (2018).

24. Holm, M. M. et al. Mature BDNF, but not proBDNF, reduces excitability of fast-spiking interneurons in mouse dentate gyrus. J. Neurosci. Off. J. Soc. Neurosci. 29, 12412–12418 (2009).

25. Wang, F. et al. RNAscope: a novel in situ RNA analysis platform for formalin-fixed, paraffin-embedded tissues. J. Mol. Diagn. JMD 14, 22–29 (2012).

26. Telley, L. et al. Dual Function of NRP1 in Axon Guidance and Subcellular Target Recognition in Cerebellum. Neuron 91, 1276–1291 (2016).

27. Bracken, B. K. & Turrigiano, G. G. Experience-dependent regulation of TrkB isoforms in rodent visual cortex. Dev. Neurobiol. 69, 267–278 (2009).

28. Bogenmann, E. et al. Generation of mice with a conditional allele for the p75(NTR) neurotrophin receptor gene. Genes. N. Y. N 2000 49, 862–869 (2011).

29. Charalampopoulos, I. et al. Genetic Dissection of Neurotrophin Signaling through the p75 Neurotrophin Receptor. Cell Rep. 2, 1563–1570 (2012).

30. Zanin, J. P., Unsain, N. & Anastasia, A. Growth factors and hormones pro-peptides: the unexpected adventures of the BDNF prodomain. J. Neurochem. 141, 330–340 (2017).

31. Rauskolb, S. et al. Global deprivation of brain-derived neurotrophic factor in the CNS reveals an area-specific requirement for dendritic growth. J. Neurosci. Off. J. Soc. Neurosci. 30, 1739–1749 (2010).

32. Dieni, S. et al. BDNF and its pro-peptide are stored in presynaptic dense core vesicles in brain neurons. J. Cell Biol. 196, 775–788 (2012).

33. Yang, J. et al. Neuronal release of proBDNF. Nat. Neurosci. 12, 113–115 (2009).

34. Yang, J. et al. proBDNF negatively regulates neuronal remodeling, synaptic transmission, and synaptic plasticity in hippocampus. Cell Rep. 7, 796–806 (2014).

35. Woo, N. H. et al. Activation of p75NTR by proBDNF facilitates hippocampal long-term depression. Nat. Neurosci. 8, 1069–1077 (2005).

36. Winnubst, J., Cheyne, J. E., Niculescu, D. & Lohmann, C. Spontaneous Activity Drives Local Synaptic Plasticity In Vivo. Neuron 87, 399–410 (2015).

37. Je, H. S. et al. ProBDNF and mature BDNF as punishment and reward signals for synapse elimination at mouse neuromuscular junctions. J. Neurosci. Off. J. Soc. Neurosci. 33, 9957–9962 (2013).

38. Pang, P. T. et al. Cleavage of proBDNF by tPA/plasmin is essential for long-term hippocampal plasticity. Science 306, 487–491 (2004).

39. Schwartz, N., Schohl, A. & Ruthazer, E. S. Activity-dependent transcription of BDNF enhances visual acuity during development. Neuron 70, 455–467 (2011).

40. Hong, E. J., McCord, A. E. & Greenberg, M. E. A biological function for the neuronal activity-dependent component of Bdnf transcription in the development of cortical inhibition. Neuron 60, 610–624 (2008).

41. Hippenmeyer, S. et al. A developmental switch in the response of DRG neurons to ETS transcription factor signaling. PLoS Biol. 3, e159 (2005).

42. Xu, Q., Tam, M. & Anderson, S. A. Fate mapping Nkx2.1-lineage cells in the mouse telencephalon. J. Comp. Neurol. 506, 16–29 (2008).

43. Pizzorusso, T. et al. Reactivation of ocular dominance plasticity in the adult visual cortex. Science 298, 1248–1251 (2002).

44. Morishita, H., Cabungcal, J.-H., Chen, Y., Do, K. Q. & Hensch, T. K. Prolonged Period of Cortical Plasticity upon Redox Dysregulation in Fast-Spiking Interneurons. Biol. Psychiatry 78, 396–402 (2015).

45. Lehmann, K. & Löwel, S. Age-dependent ocular dominance plasticity in adult mice. PloS One 3, e3120 (2008).

46. Naska, S., Lin, D. C., Miller, F. D. & Kaplan, D. R. p75NTR is an obligate signaling receptor required for cues that cause sympathetic neuron growth cone collapse. Mol. Cell. Neurosci. 45, 108–120 (2010).

47. Sun, Y. et al. ProBDNF collapses neurite outgrowth of primary neurons by activating RhoA. PloS One 7, e35883 (2012).

48. Deinhardt, K. et al. Neuronal growth cone retraction relies on proneurotrophin receptor signaling through Rac. Sci. Signal. 4, ra82 (2011).

49. Yamashita, T. & Tohyama, M. The p75 receptor acts as a displacement factor that releases Rho from Rho-GDI. Nat. Neurosci. 6, 461–467 (2003).

50. Yamashita, T., Fujitani, M., Yamagishi, S., Hata, K. & Mimura, F. Multiple signals regulate axon regeneration through the Nogo receptor complex. Mol. Neurobiol. 32, 105–111 (2005).

51. Lim, Y.-S. et al. p75(NTR) mediates ephrin-A reverse signaling required for axon repulsion and mapping. Neuron 59, 746–758 (2008).

52. Baho, E. & Di Cristo, G. Neural activity and neurotransmission regulate the maturation of the innervation field of cortical GABAergic interneurons in an age-dependent manner. J. Neurosci. Off. J. Soc. Neurosci. 32, 911–918 (2012).

53. Gao, X., Daugherty, R. L. & Tourtellotte, W. G. Regulation of low affinity neurotrophin receptor (p75NTR) by early growth response (Egr) transcriptional regulators. Mol. Cell. Neurosci. 36, 501–514 (2007).

54. Irmady, K. et al. Mir-592 regulates the induction and cell death-promoting activity of p75NTR in neuronal ischemic injury. J. Neurosci. Off. J. Soc. Neurosci. 34, 3419–3428 (2014).

55. Unsain, N., Nuñez, N., Anastasía, A. & Mascó, D. H. Status epilepticus induces a TrkB to p75 neurotrophin receptor switch and increases brain-derived neurotrophic factor interaction with p75 neurotrophin receptor: an initial event in neuronal injury induction. Neuroscience 154, 978–993 (2008).

56. Volosin, M. et al. Induction of proneurotrophins and activation of p75NTR-mediated apoptosis via neurotrophin receptor-interacting factor in hippocampal neurons after seizures. J. Neurosci. Off. J. Soc. Neurosci. 28, 9870–9879 (2008).

57. Bavelier, D., Levi, D. M., Li, R. W., Dan, Y. & Hensch, T. K. Removing brakes on adult brain plasticity: from molecular to behavioral interventions. J. Neurosci. Off. J. Soc. Neurosci. 30, 14964–14971 (2010).

58. Carulli, D. et al. Animals lacking link protein have attenuated perineuronal nets and persistent plasticity. Brain J. Neurol. 133, 2331–2347 (2010).

59. Nykjaer, A. et al. Sortilin is essential for proNGF-induced neuronal cell death. Nature 427, 843–848 (2004).

60. Teng, H. K. et al. ProBDNF induces neuronal apoptosis via activation of a receptor complex of p75NTR and sortilin. J. Neurosci. Off. J. Soc. Neurosci. 25, 5455–5463 (2005).

61. Kolbeck, R., Jungbluth, S. & Barde, Y. A. Characterisation of neurotrophin dimers and monomers. Eur. J. Biochem. 225, 995–1003 (1994).

62. Mizui, T. et al. BDNF pro-peptide actions facilitate hippocampal LTD and are altered by the common BDNF polymorphism Val66Met. Proc. Natl. Acad. Sci. U. S. A. 112, E3067–3074 (2015).

63. Chen, Z.-Y. et al. Genetic variant BDNF (Val66Met) polymorphism alters anxiety-related behavior. Science 314, 140–143 (2006).

64. Soliman, F. et al. A genetic variant BDNF polymorphism alters extinction learning in both mouse and human. Science 327, 863–866 (2010).

65. Zhang, L. et al. PTSD risk is associated with BDNF Val66Met and BDNF overexpression. Mol. Psychiatry 19, 8–10 (2014).

66. Porciatti, V., Pizzorusso, T. & Maffei, L. The visual physiology of the wild type mouse determined with pattern VEPs. Vision Res. 39, 3071–3081 (1999).

67. Groleau, M. et al. Impaired functional organization in the visual cortex of muscarinic receptor knock-out mice. NeuroImage 98, 233–242 (2014).

